# Pyroptolysins are versatile, sterol-sensing bacterial toxins

**DOI:** 10.64898/2026.10.05.756811

**Authors:** MacKenzie Patterson, Anna L. Rakowski, Emma Kote, Angelina C. Como-Mosconi, Bin Huang, Megan L. Mayer, Alex G. Johnson

## Abstract

Gasdermin pores mediate cellular self-sacrifice for the immune defense pyroptosis. Here we describe how the gasdermin fold was repurposed during evolution into a membrane targeting bacterial toxin. These proteins, which we call pyroptolysins, are natural chimeras of gasdermins and cholesterol-dependent cytolysins. Like the cytolysins, pyroptolysins are active on cholesterol-rich membranes, but unlike the cytolysins, they are poorly active on mammalian cells. Structural snapshots of the pyroptolysins confirm their dual homology and reveal pre-organized complexes primed for membrane insertion. Unexpectedly, we found the pyroptolysins to be active *in vitro* and *in vivo* on membranes with remarkably low sterol content, displaying insecticidal activity. Our results establish the gasdermin fold as a versatile pore-forming scaffold, capable of inducing cell death from the inside and the outside of cells.

---

In animal, fungal, and bacterial immunity, gasdermins form membrane pores that induce host cell suicide following signaling events that result from nonself recognition^1–6^. Gasdermins are typically activated by proteolysis of an autoinhibitory C-terminal domain, though cleavage-independent mechanisms have recently been uncovered in animals and fungi^7,8^. In contrast to the superfamily of complement <u>m</u>embrane <u>a</u>ttack <u>c</u>omplex component, <u>p</u>erforin, and <u>c</u>holesterol-<u>d</u>ependent <u>c</u>ytolysin (MACPF/CDC) pore-forming proteins^9–11^, gasdermins invariably act from within cells, where they typically target the inner leaflet of the plasma membrane. Despite undetectable sequence homology, the gasdermins and MACPF/CDC proteins assemble into structurally similar β-barrel pores, with each protomer contributing four β strands to ∼16–52 subunit assemblies^9,11,12^. While formerly considered to have arisen via convergent evolution^13^, a preliminary query of the Alphafold database identified bacterial proteins that are apparent chimeras of both gasdermins and CDCs^14^. However, the membrane targets, activation mechanism, and physiological role of these proteins remained unknown.

Here, we name the gasdermin–CDC chimeric proteins “pyroptolysins”. To reveal their natural diversity, we exhaustively searched sequence databases and built a phylogenetic tree of non-redundant sequences (Figure 1 A–B; Figure S1A). Pyroptolysins are encoded by bacteria and archaea, but in contrast to the CDCs which are primarily encoded by gram-positive pathogens^15^, the pyroptolysins are enriched in environmental, gram-negative bacteria (Figure 1B; Figure S1B). To confirm their predicted homology to gasdermins and CDCs, we solved a crystal structure of a pyroptolysin from *Pseudomonas putida* (Table S1). The *Pseudomonas* pyroptolysin comprises a gasdermin-like N-terminal domain in its resting, inactive state, and a fused CDC-like β- sandwich C-terminal domain (Figure 1C). The N-terminal domains of *Pseudomonas* pyroptolysin and *Vitiosangium* gasdermin^6^ share close structural homology in the palm, wrist, and thumb domains, but the pyroptolysin lacks certain α-helices in its proto-finger region (Figure S2A–B), while the C-terminal domain closely resembles the Ig-like membrane targeting domain of pneumolysin (Figure S2C).

**Figure 1.**
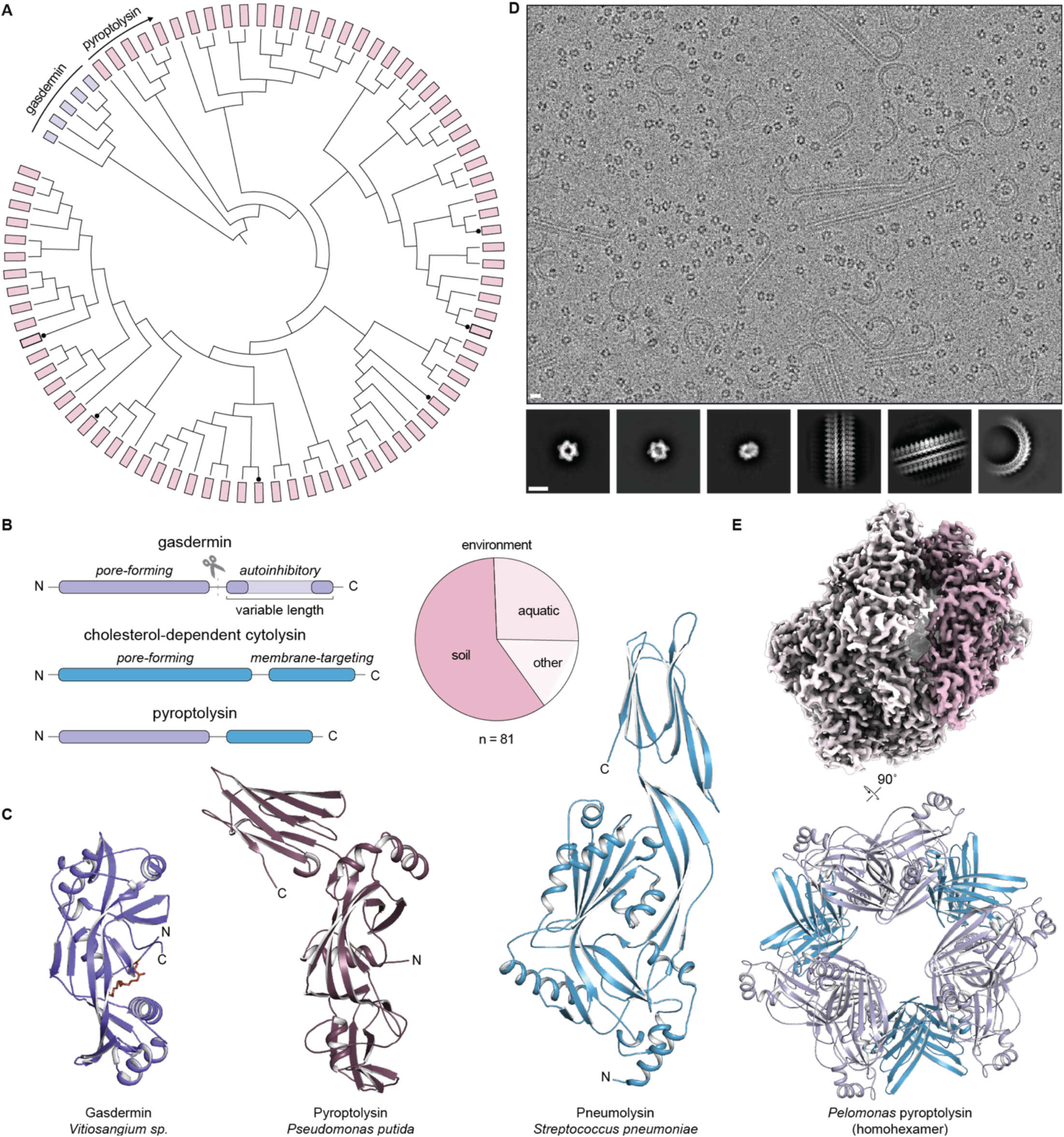
Pyroptolysins are natural chimeras of gasdermins and cholesterol-dependent cytolysins. (A) Phylogenetic tree of pyroptolysins. Node tips bars are sized by sequence length, tips labeled with a “•” are proteins characterized in this study, and tip bars outlined in black are the *Pseudomonas* and *Pelomonas* pyroptolysins. Bacterial gasdermins were used as an outgroup. (B) Left, domain architecture diagrams of gasdermins, pyroptolysins, and cholesterol-dependent cytolysins. Right, pie chart of environments occupied by pyroptolysin-encoding bacteria from the tree in panel A. (C) Crystal structures of a bacterial gasdermin (Protein Data Bank ID: 7N51), the *Pseudomonas putida* pyroptolysin, and pneumolysin (Protein Data Bank ID 5AOD). (D) Cryo-EM micrograph and 2D classes of *Pelomonas* pyroptolysin. Scale bar, 10 nm. Cryo-EM micrograph and 2D classes of *Pelomonas* pyroptolysin. Scale bar, 10 nm. (E) Legends should refer to relevant items in the supplemental information (e.g., “See also Figure S1.”).

Early cryo-electron microscopy studies of pneumolysin revealed its tendency to oligomerize into helical assemblies in aqueous solution^16,17^. While screening pyroptolysins for crystallography, we found that some homologs spontaneously assembled into filaments, in agreement with their homology to CDCs (Figure 1D; Figure S1D). An exceptional homolog from a *Pelomonas* species was found to form “stars and stripes”-like assemblies, which allowed us to determine a 2.4 Å-resolution cryo-EM structure of a homohexameric particle with *D_3_* symmetry (Figure 1E; Figure S3; Table S2). A model of the *Pelomonas* pyroptolysin confirmed an inactive-state conformation of the gasdermin N-terminal domain, with inter-subunit interactions stabilized by an extensive interface, mediated in part by a unique N-terminal extension that β-augments a kinked β3-strand in the proto-finger region (Figure S3).

Inspecting their genomic neighborhood revealed that pyroptolysins are primarily encoded as solitary genes (Figure 2A), in stark contrast to the proteolytically activated bacterial gasdermins^6^. Interestingly, one pyroptolysin from *Pseudomonas alcaligenes* encoded a neighboring protein with an ADP-ribosyltransferase (ART) fold with structural homology to a *Pseudomonas syringae* pathogenic effector (Figure S4A, DALI Z-score = 5.8), suggesting that pyroptolysins might function in interspecies antagonism^18–21^. We therefore hypothesized that pyroptolysins intoxicate non-mammalian organisms inhabiting their ecological niches. To first test this hypothesis for bacteria, we expressed four diverse pyroptolysins from within *E. coli* cells but observed no evidence for toxicity (Figure S4B). To test this hypothesis for a model eukaryote, we integrated pyroptolysins into the *S. cerevisiae* genome under control of a galactose-inducible promoter. In contrast to the *E. coli* experiments, overexpressing pyroptolysins from yeast cells potently inhibited growth, suggesting that pyroptolysins might target eukaryotes with sterol-containing membranes (Figure 2B; Figure S4C). However, external addition of pyroptolysins to growing yeast cells caused no apparent effect to growth rate, possibly due to their inability to penetrate the fungal cell wall (Figure S4D).

**Figure 2.**
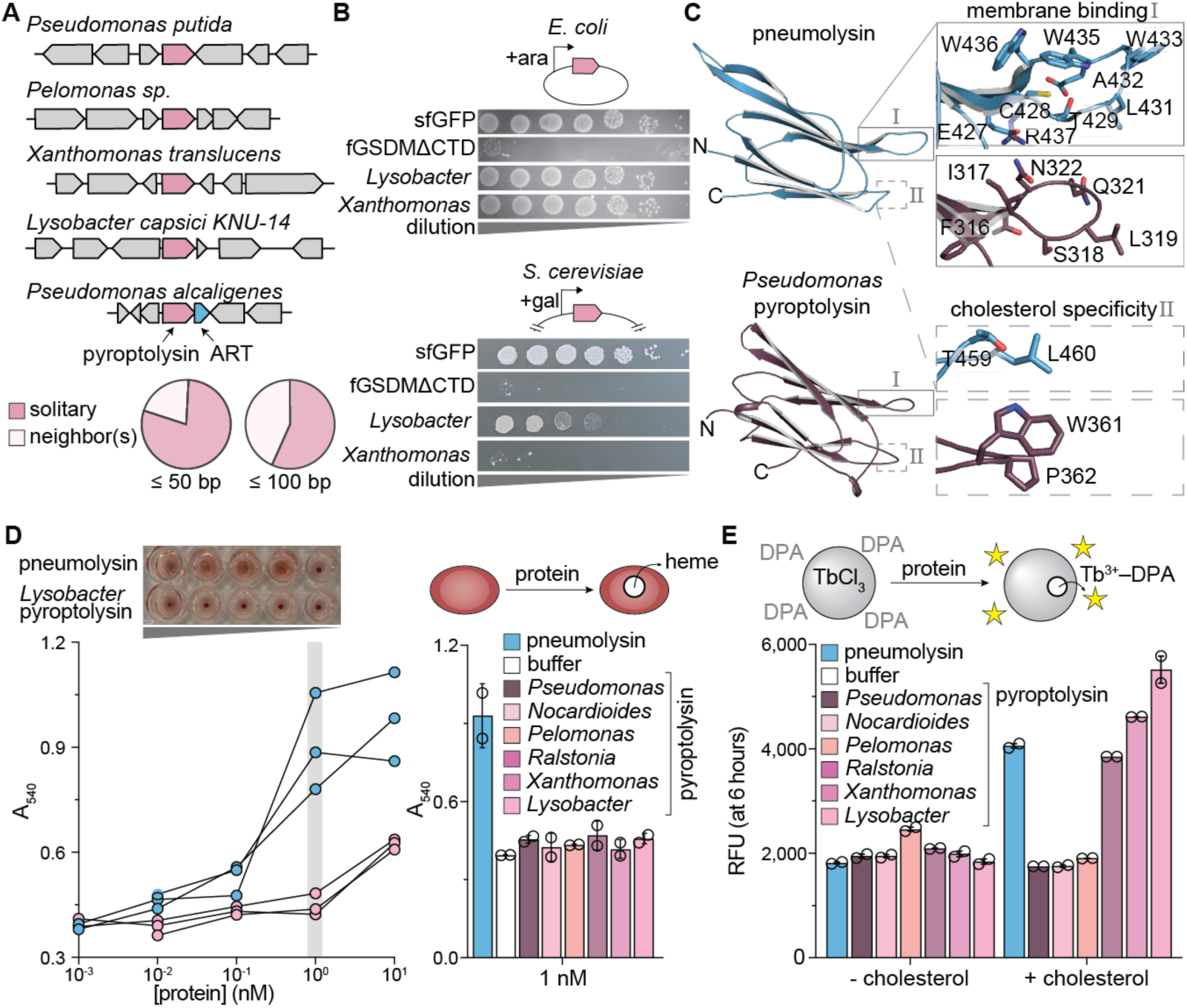
Pyroptolysins are active on sterol-containing membranes but spare mammalian cells. (A) Neighborhood diagrams of select pyroptolysin genes (top). An example neighbor encoding a putative ADP-ribosyltransferase (ART) is highlighted in blue. Pie charts show species categorized based on whether they have a neighboring gene encoded on the same strand within the denoted cutoff (bottom, n = 370 sequences). (B) Spot assays of *E. coli* (top) or *S. cerevisiae* (bottom) cells induced to express genes encoding sfGFP, an autoactive fungal gasdermin, or pyroptolysins from inducible promoters. (C) Comparison of membrane-binding motifs in the Ig-like domains from pneumolysin (Protein Data Bank ID 5AOD) and the *Pseudomonas* pyroptolysin. Insets show motifs for CDC membrane binding (the undecapeptide) and cholesterol recognition (cholesterol specificity) and equivalent positions on the *Pseudomonas* pyroptolysin structure. (D) Hemolysis assay of pneumolysin and pyroptolysins on sheep erythrocytes. A representative image of erythrocytes after incubation with varying concentrations of proteins is shown above a dose-response curve of hemolysis for select proteins (left, gray bar = 1 nM). Hemolytic activity of 1 nM proteins is shown on the right. (E) Relative fluorescent units (RFUs) at six hours from a liposome rupture experiment testing proteins on DOPC liposomes reconstituted with 25 mol% cholesterol.

Given the potential activity of pyroptolysins on sterol-containing membranes, we compared the Ig-like domains of pyroptolysin and CDC by structure-based sequence alignments. While CDCs contain a tryptophan-rich “undecapeptide” loop that is important for membrane anchoring^15^, the pyroptolysins have an abridged loop, with little sequence conservation between homologs (Figure 2C; Figure S2C). Additionally, CDCs bind cholesterol with a conserved “cholesterol specificity” threonine-leucine (TL) pair^22^, while pyroptolysins share a highly conserved tryptophan-proline (WP) motif in its place (Figure 2C, Figure S2C). Despite these differences, we asked whether the pyroptolysins might target mammalian cells using hemolytic assays. Screening a panel of six diverse pyroptolysins, we observed very weak hemolytic activity in comparison to the CDC pneumolysin (Figure 2D; Figure S4E) and additionally confirmed weak cytotoxicity to human HEK293 cells (Figure S4F). Despite their lack of activity on naturally cholesterol-rich membranes, we asked whether membrane cholesterol could activate pyroptolysins *in vitro*. Using 1,2-dioleoyl-sn-glycero-3-phosphocholine (DOPC) liposomes containing cholesterol at a 25% molar ratio, we screened the same panel of six pyroptolysins using liposome rupture assays. Intriguingly, most pyroptolysins ruptured these liposomes, some to a greater extent than pneumolysin (Figure 2E). Together with their potent cell killing of yeast cells, the *in vitro* activity of pyroptolysins on cholesterol-containing liposomes suggested that the proteins target eukaryotic sterol-containing membranes, but do not naturally target mammalian cells.

To investigate the structural basis for the pyroptolysin pore-forming activity, we incubated pyroptolysins with DOPC liposomes containing 25 mol% cholesterol or ergosterol, being the fungal sterol equivalent. By negative-stain electron microscopy, pores were found to stud liposomes with inner diameters ranging from ∼20– 30 nm, independent of sterol type (Figures S4F and S4G. We prepared *Lysobacter* pyroptolysin pore–liposome samples for cryo-electron microscopy and resolved single pores and head-to-head pore dimers bridging sterol-containing liposomes (Figure 3A). A protein and detergent screen identified nonylphenyl polyethylene glycol as a favorable detergent for extracting the *Lysobacter* pyroptolysin pores from the liposomes. After exchanging extracted pores into amphipols, this sample was used to determine a 2.8 Å-resolution cryo-EM structure of the 39-subunit *Lysobacter* pyroptolysin pore as a head-to-head dimer (Figure 3B; Figure S5A, Table S2). The *Lysobacter* pyroptolysin pore adopted a similar oligomerization stoichiometry and dimensions to the 42-mer pneumolysin pore^23^ with an inner diameter of 250 Å, height of 100 Å, and outer diameter of 350 Å (Figures 3B and 3C). The high-resolution structure enabled atomic model building of a complete active-state *Lysobacter* pyroptolysin, which adopted a gasdermin “hand shaped” fold with the CDC Ig-like domain lining the outside of the pore at an extended angle with respect to pneumolysin (Figure 3D; Figure S5B).

**Figure 3.**
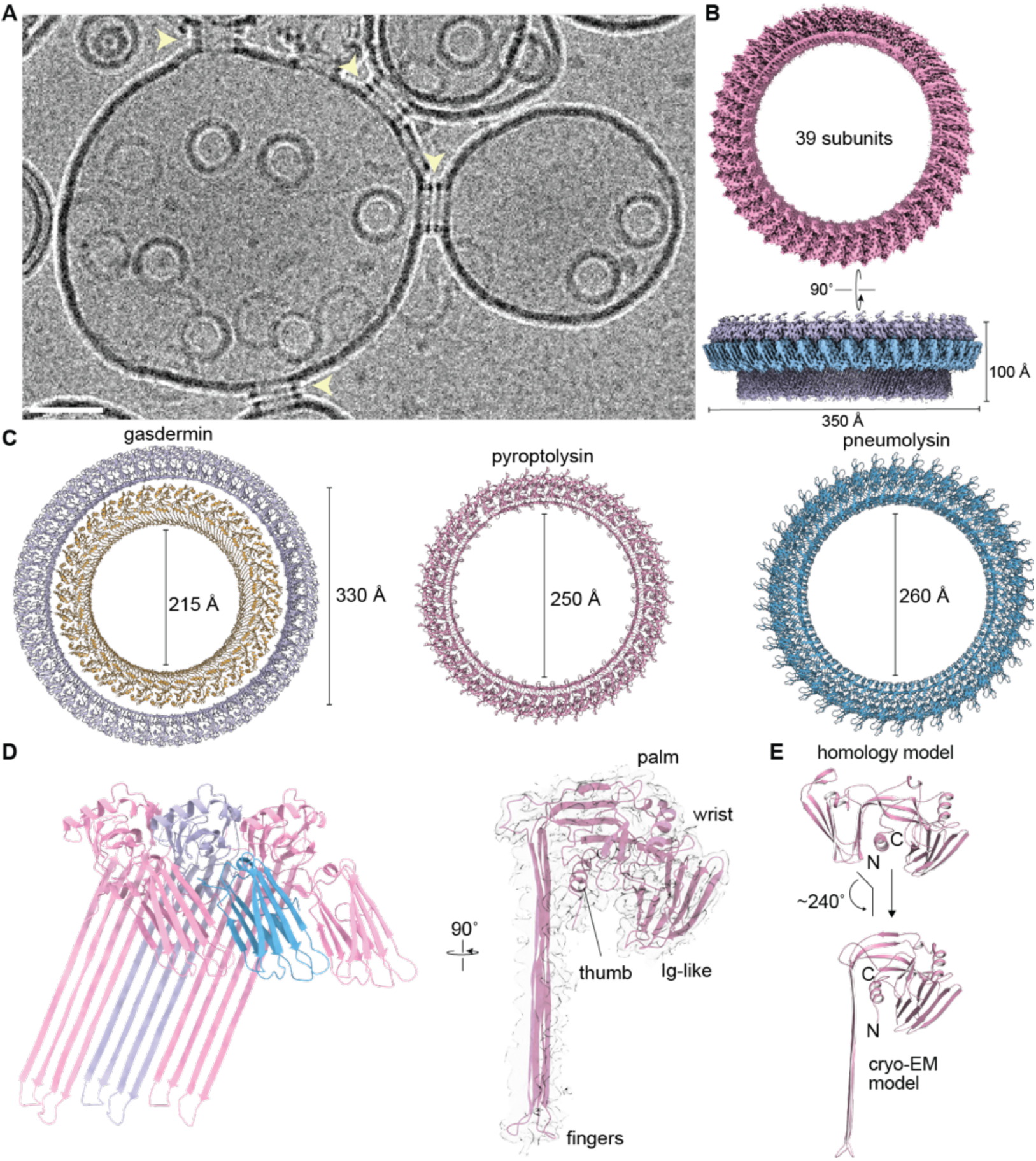
Structure of a pyroptolysin membrane pore. (A) A cryo-electron micrograph of *Lysobacter* pyroptolysin pores on DOPC liposomes with 25 mol% ergosterol. Yellow arrows indicate stacked pores. Scale bar, 50 nm. (B) 2.8 Å-resolution cryo-EM map of the *Lysobacter* pyroptolysin pore extracted from DOPC liposomes with 25 mol% ergosterol. (C) Comparative dimensions of *Vitiosangium* bacterial gasdermin (purple, Protein Data Bank ID 8SL0), human gasdermin D (yellow, Protein Data Bank ID 6VFE), *Lysobacter* pyroptolysin, and pneumolysin (Protein Data Bank ID 5LY6) pores. (D) Map-model overlay of a *Lysobacter* pyroptolysin protomer modeled into the map shown in panel B. (E) The structural transition for activation of a *Lysobacter* pyroptolysin protomer.

Comparing a homology model of the inactive *Lysobacter* pyroptolysin with the structure of the active protomer revealed a ∼240° unfurling of proto-finger β-strands into the transmembrane β-sheet, but minimal secondary structure changes with a 5% α–β transition in the finger regions compared to the more extensive 23% for gasdermin and 31% for CDC activation mechanisms (Figure 3E; Figure S6A; Table S3). We therefore hypothesized that pyroptolysins are uniquely poised for membrane insertion and might exhibit a distinct response to sterol recognition.

To probe their sterol specificity, we screened four natural membrane sterols in liposome rupture assays, finding that pyroptolysins ruptured liposomes with 25% mol% of all sterols tested and in nearly all cases showing enhanced activity versus pneumolysin (Figure 4A). This lack of specificity prompted us to examine whether pyroptolysin activation was sterol-dependent or whether they instead respond to sterol-induced changes in membrane fluidity. We therefore tested pyroptolysins against liposomes containing 3-epicholesterol, an unnatural epimer of cholesterol with the C3 hydroxyl in the α-configuration instead of the natural β-configuration^24^. Supporting the requirement for direct sterol recognition, pyroptolysins failed to rupture these liposomes. We next tested whether modulating membrane sterol accessibility might affect pyroptolysin activity. Pyroptolysins showed reduced rupture of palmitoyl-2-oleoyl-glycero-3-phosphocholine (POPC) versus DOPC liposomes, consistent with tighter lipid packing due to one less unsaturation on the POPC acyl chain with respect to DOPC (Figure S6B)^24^. Similarly, pyroptolysins also showed higher activity towards DOPC liposomes containing 25% cholesterol and 10% sphingomyelin, known to increase sterol accessibility via formation of lipid rafts (Figure S6C)^25^.

**Figure 4.**
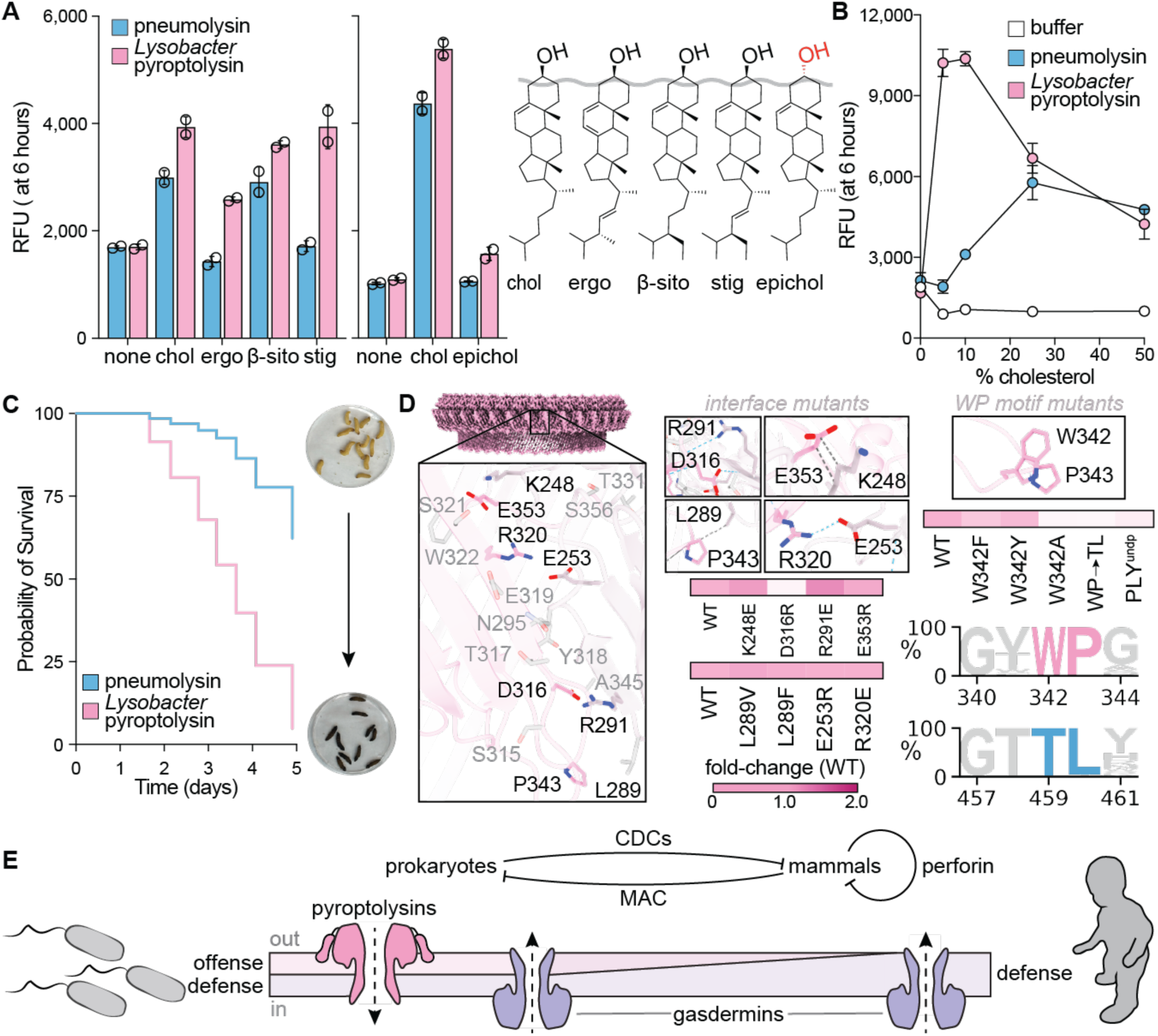
Pyroptolysins target sterol-containing liposomes with unprecedented sensitivity. (A) Liposome rupture experiment testing pneumolysin and *Lysobacter* pyroptolysin on DOPC liposomes with 25 mol% sterol (left to right, cholesterol, ergosterol, β-sitosterol, stigmasterol, and epicholesterol). (B) Liposome rupture experiment testing pneumolysin and pyroptolysins on DOPC liposomes reconstituted with 5, 10, 25, or 50 mol% cholesterol. For panel A–B, Relative fluorescent units (RFUs) are measured six hours following protein addition. (C) Kaplan-Meier survival curves of *Galleria mellonella* larvae after injection with 50 ng of pneumolysin or *Lysobacter* pyroptolysin. Image of live worms is from a PBS control. (D) Left, Ig-like domain interface of neighboring active-state *Lysobacter* pyroptolysin subunits in the pore. Right, normalized heat maps (relative to WT *Lysobacter* pyroptolysin) for liposome rupture activity of mutants against DOPC liposomes with 25 mol% cholesterol. Insets to the right show the mutations on the active-state *Lysobacter* pyroptolysin structure in stick representation. Bottom right, sequence logos of the conserved “WP” and “TL” motifs in pyroptolysins and CDCs. (E) A model for the evolution of the β-barrel pore-forming proteins in pathogen offense and defense.

These results suggested that sterol accessibility altered the efficiency of pyroptolysin pore formation. We therefore began titrating the levels of cholesterol in our experiments. Multiple pyroptolysins ruptured POPC liposomes with 25 mol% cholesterol, which is below the threshold typically required to fully activate CDCs *in vitro* (Figure S6D)^26–28^. Both the *Pseudomonas* and *Pelomonas* pyroptolysins showed poor activity even on 50% cholesterol containing liposomes, suggesting they might be activated by alternative mechanisms. We hypothesized that removal of the unusual *Pelomonas* pyroptolysin N-terminal extension might function as an autoinhibitory domain via interactions across the oligomerization interface and its removal would induce pore-formation. We first attempted to delete this extension (residues 1–21) but found that its removal prevented purification of a soluble protein. Inspired by studies of CDC-like proteins that used broad-specificity proteases to activate pore formation when the cognate protease is unknown^29^, we tested proteinase K activation of the *Pelomonas* pyroptolysin. Excitingly, we found that proteinase K strongly stimulated rupture activity towards liposomes with but not without cholesterol, supporting a proteolytic, sterol-dependent mechanism for some pyroptolysins (Figure S6E).

Remarkably, pyroptolysins ruptured DOPC liposomal membranes with cholesterol concentrations as low as 5% (Figure 4B, Figure S6D). The *Lysobacter* pyroptolysin exhibited optimal activity in DOPC liposomes with 5–10% cholesterol, representing a significant shift in sensitivity with respect to CDCs. Desulfolysin, a gram-negative CDC encoded by bacteria that are not known to be human pathogens, was also found to exhibit cholesterol sensitivity nearly on par with the *Lysobacter* pyroptolysin (Figure S6F). Given this surprising activity, we hypothesized that pyroptolysins might target cell types with naturally low membrane sterol content. To test this hypothesis, we applied pyroptolysins exogenously to *Spodoptera frugiperda* (Sf9) insect cells, which have <4 mol% cholesterol in their membranes^30^. Relative to pneumolysin, pyroptolysins showed enhanced cytotoxicity to Sf9 cells, a complete reversal of our previous hemolysis assay results (Figure S7A). Extending this hypothesis to an insect animal model, we observed dose-dependent toxicity against the wax worm *Galleria mellonella* (Figure 4C, Figure S7), which was again enhanced for pyroptolysins versus pneumolysin.

To probe the molecular features in the Ig-like domain that influence pyroptolysin membrane targeting and oligomerization, we examined the *Lysobacter* pyroptolysin pore model. Adjacent Ig-like β-sandwiches form an extensive zipperlike interface distinct from the pneumolysin pore (Figure 4D, Figure S8)^23^. Most charge swap mutations failed to perturb liposome rupture activity, but alanine substitution of the conserved tryptophan in the “WP” motif led to strongly diminished activity (Figure 4D, Figure S8). Substituting the “WP” motif for the CDC “TL” motif similarly diminished pyroptolysin activity, but substitution of the tryptophan with aromatic residues had little effect. Swapping in the CDC undecapeptide also significantly reduced activity, indicating that the pyroptolysins follow a distinct sterol recognition mechanism from CDCs (Figure 4D, Figure S8). The “WP” motif potentially contributes to this mechanism by forming stronger Ch–π interactions with the sterol backbone versus the transient interactions of CDCs^31^.

Our discovery and mechanistic analysis of the pyroptolysins clarifies the evolutionary link between the gasdermins and the MACPF/CDC superfamily, revealing the versatile nature of the gasdermin fold in pathogen defense and offense (Figure 4E)^13^. Despite their structural homology to CDCs and their activity on sterol-containing membranes, pyroptolysins do not target mammalian cells and use a dissimilar sequence motif to target sterol-containing membranes. Notably, the gram-negative CDC desulfolysin also exhibits higher sterol sensitivity than pneumolysin and perfringolysin O (Figure S6F), but unlike the pyroptolysins it contains CDC-like motifs and is hemolytically active^32,33^. Membrane sterols are therefore necessary but not sufficient for pyroptolysin cellular activity. Sphingomyelin stimulated the cholesterol-dependent rupture activity of some pyroptolysins (Figure S6C), suggesting that pyroptolysins might recognize sterol-lipid complexes like the fungal toxin OlyA/PlyB^34^. Given that CDCs recognize glycan and protein receptors^35–37^, we propose that pyroptolysins may similarly bind non-lipid receptors to target organisms within their ecological niche, including insect cells. Such intriguing behavior is akin to the recently described insecticidal activity of homologs of the classical botulinum and diptheria toxins^38,39^. While rare, the existence of two-gene pyroptolysins/ART systems suggests that some pyroptolysins may be involved in delivery of toxin payloads^40^, further highlighting their versatility for different targeting mechanisms. Engineering the pyroptolysin scaffold for selective cell targeting could provide an opportunity to exogenously induce pyroptosis for host defense, while sparing healthy cells. Further work aimed at deconvoluting the environmental roles and organismal targets of pyroptolysins will both explain the evolutionary history of gasdermins in pathogen offense, illuminate the molecular principles that guide sterol recognition, and enable the design of membrane toxins with useful functions.

## Acknowledgements

The authors thank members of their lab, A. Heuck, M. Burroughs, H. Ludewig, K. Ragunathan, and J. Haber for helpful discussions. We thank A. Latuda for assistance with yeast experiments. TEM samples were prepared and imaged at the Brandeis Electron Microscopy Facility, which made use of equipment purchased under National Science Foundation MRI Program Award #2319804. We thank A. Tiano for her assistance with the F20 microscope. Cryo-EM data was collected at the Harvard Cryo-EM Center for Structural Biology at Harvard Medical School and at the PNCC supported by NIH grant U24GM129547. We thank O. Davulcu for expert assistance with electron microscopy.

## Author Contributions

Conceptualization: MP, AGJ

Methodology: MP, EK, AR, AGJ

Investigation: MP, EK, AR, AC-M, BH, AGJ

Visualization: MP, AGJ

Funding acquisition: AGJ

Project administration: AGJ

Supervision: AGJ

Writing – original draft: MP, AGJ

Writing – review & editing: MP, AGJ

## Competing Interests

MP, EK, and AGJ are inventors on a provisional patent application filed by Brandeis University on the work described in this study.

## Materials and Methods

### Phylogenetic analysis of pyroptolysin sequences

Pyroptolysin sequences were obtained from the AlphaFold^42^ database using experimentally determined bacterial gasdermin sequences as queries^14^. The following pyroptolysin sequences were used as queries against the NCBI^43^ protein BLAST database: *Pseudomonas* (IMG Gene ID: 2637766654), *Lysobacter* (IMG Gene ID: 8123842218), *Xanthomonas* (IMG Gene ID: 2819659989) and *Ralstonia* (IMG Gene ID: 2859182761). The nonredundant set of sequences was clustered using mmseqs2^44^ version 18.8cc5c by 80% sequence identity with a minimum target coverage threshold of 80% (settings –min-seq-id 0.8 –c 0.8 –cov-mode 1). The four query pyroptolysins were prioritized as the representative members of their clusters so they would be included in the final list of sequences. Clustered sequences were filtered for length between 325 and 400 amino acids to remove bacterial gasdermins, CDCs, and solitary Ig-like domains. A multiple sequence alignment was generated from the set of filtered sequences using Clustal Omega^45^ version 1.2.3 using five bacterial gasdermin sequences as an outgroup (NCBI references numbers: WP_108071778.1, WP_157872517.1, WP_203409785.1, WP_277336092.1, and WP_266940482.1). A phylogenetic tree was calculated from the multiple sequence alignment using FastTree^46^ version 2.1.11 employing the Jones-Taylor-Thornton evolutionary model and visualized and annotated in iTOL^47^. In addition to the pyroptolysin sequences used as queries, *Nocardioides* (IMG Gene ID: 2996130405), and *Pelomonas* (IMG Gene ID: 2643742627) pyroptolysins were selected for further study.

Taxonomy and gene neighborhood analysis were performed using a pipeline executed by Claude code (Anthropic, accessed August 2026). The pipeline used NCBI Entrez^43^ efetch for each pyroptolysin sequence against the identical protein groups (IPG) resource to retrieve its nucleotide accession number, start and stop indices, strand sense, source organism, and assembly identifier. The pipeline then extracted 20 kb on either side of the pyroptolysin start/stop indices and used Bio.SeqIO^48^ to return a GenBank record. For each neighboring CDS, the pipeline retrieved its start and stop indices, strand, NCBI protein accession number, gene product, locus tag, gene and translation. Coordinate distances were measured between the pyroptolysin CDS and any neighbors present within 50 bp or 100 bp.

Neighbors with “hypothetical protein” or “uncharacterized” product names were submitted to the EBI InterProScan^49^ web service in three batches ≤ 100 sequences each with all default analyses enabled. Returned JSON files were parsed to extract the best Pfam^50^ hit, InterPro accessions^49^, transmembrane helix count, and signal peptide flags. Neighbors that did not have a Pfam/InterPro assignment were submitted to AlphaFold 3^51^. Sequences with a predicted signal peptide were submitted with residues 1-25 trimmed to prevent downstream confusion about fold comparison. For each neighbor, the 60 top-ranked models were searched with Foldseek^52^ v. 10 using easy-search with maximum sensitivity (-s 9.5) against PDB100, AlphaFold DB Swiss-Prot, and CATH50 databases.

Sequence logos of pyroptolysins and CDCs were created using Logomaker^53^. A CDC sequence alignment was generated by querying pneumolysin against the NCBI^43^ protein BLAST database, filtering for length between 400 and 600 amino acids, and aligning using Clustal Omega^45^ version 1.2.3.

### Recombinant protein expression and purification

Pyroptolysin genes were codon optimized for expression in *E. coli* and synthesized and cloned into a custom pET expression vector encoding an N-terminal 6×His-tagged human SUMO2 tag (pETS2) by Twist Biosciences. The N-terminal methionine residue was removed from each sequence and replaced by a single serine residue following hSUMO2 tag removal by the human SENP2 protease. Mutants were prepared by site-directed mutagenesis using custom primers (Genewiz) or DNA synthesis (Twist Biosciences) and assembled using Gibson Assembly into pETS2.

For expression, BL21(DE3)-RIL *E. coli* cells were transformed with pETS2 plasmids and plated on LB agar plates containing 100 mg mL^−1^ ampicillin, 34 mg mL^−1^ chloramphenicol and 1% glucose. One to five of the resulting colonies were used to inoculate starter cultures of MDG medium for overnight growth at 37 °C. MDG (0.5% glucose, 25 mM Na_2_HPO_4_, 25 mM KH_2_PO_4_, 50 mM NH_4_Cl, 5 mM Na_2_SO_4_, 2 mM MgSO_4_, 0.25% aspartic acid, 100 mg mL^−1^ ampicillin, 34 mg mL^−1^ chloramphenicol, and trace metals) starter cultures were used to inoculate 2×1 L of M9ZB medium (0.5% glycerol, 1% Cas-amino acids, 47.8 mM Na_2_HPO_4_, 22 mM KH_2_PO_4_, 18.7 mM NH_4_Cl, 85.6 mM NaCl, 2 mM MgSO_4_, 100 mg mL^−1^ ampicillin, 34 mg mL^−1^ chloramphenicol, and trace metals) and grown at 37 °C with shaking at 180 rpm until the culture reached an OD_600_ of ∼2.5. Cultures were chilled on ice for ∼20 min and induced for expression with 0.5 mM IPTG before shaking overnight at 16 °C and 180 rpm. The next morning, expression cultures were pelleted by centrifugation for 15 min at 4,000 *×g*, flash frozen on liquid nitrogen and stored at −80 °C until purification.

Protein purification was performed at 4 °C with all buffers containing 20 mM HEPES-KOH (pH 7.5) and other components as described below. Cell pellets were thawed and lysed by sonication in buffer containing 400 mM NaCl, 30 mM imidazole, with or without 1 mM DTT. Lysate was clarified by centrifugation and passage through glass wool, bound to NiNTA agarose beads (QIAGEN, ThermoScientific), which were subsequently washed with buffer containing 1 M NaCl, 30 mM imidazole, and DTT when used. Protein was eluted with buffer containing 400 mM NaCl, 300 mM imidazole, and DTT when used. Human SENP2 protease was added to the eluate to cleave off the SUMO2 tag before dialyzing overnight into buffer containing 250 mM KCl with or without 1 mM DTT. For initial screening of candidates, the dialyzed protein was directly purified by size-exclusion chromatography (SEC) using a 16/600 Superdex 75 or 16/600 Superdex 200 column (Cytiva) equilibrated with 250 mM KCl and 1 mM Tris(2-carboxyethyl)phosphine hydrochloride (TCEP). Sized proteins were brought to concentrations >20 mg mL^−1^, using 10 kDa molecular weight cut-off concentrators (Millipore), flash frozen on liquid nitrogen, and stored at −80 °C until use.

### Protein crystallization and structure determination

Proteins for crystallization were thawed from –80 °C stocks on ice and diluted to concentrations of 15 mg mL^-1^ protein in 20 mM HEPES-KOH (pH 7.5). Crystallization conditions were screened using the hanging-drop vapor diffusion method with a SPT Labtech Mosquito crystallization robot and Hampton screens. Crystallization conditions for *Pseudomonas* pyroptolysin (0.1 M Bis-Tris, pH 6.5, 20% w/v PEG 3,350) were optimized in 24-well VDX hanging-drop vapor diffusion trays with sealant (Hampton) with 2 µL drop volume at 2:3, 3:2, and 1:1 protein:reservoir ratios over 500 µL of reservoir. Crystals were grown for at least 1 week at 16 °C, incubated in reservoir solution supplemented with cryoprotectant (35% w/v ethylene glycol), and harvested by flash freezing on liquid nitrogen. X-ray diffraction data were acquired using Northeastern Collaborative Access Team beamline 24-ID-E (GM124165) and used an Eiger detector (OD021527) at the APS (DE-AC02-06CH11357, DOI: https://doi.org/10.46936/APS-195197/60017694). Data were processed with RAPD2 and Phenix version 1.20.1. Atomic models were built in Coot and refined in Phenix. Statistics were analyzed as described in Table S1. Structure Figures were generated using PyMOL version 2.5.0 open-source build^54^ and ChimeraX version 1.11^55^.

### Detergent screens and extraction of pyroptolysin pores

To assemble pores in liposomes, reactions of pyroptolysins (20 µM) and liposomes (∼1.0 mg mL^−1^) were incubated overnight at room temperature in glass vials. Screens were initially performed using liposome compositions found to be most ideal by negative-stain electron microscopy analysis. Reactions were carried out at room temperature overnight. The next day, liposomes were collected by centrifuging 150 µL portions at 21,000 ×*g* at 4 °C for 30 min with an Eppendorf 5425R mini centrifuge. The supernatant containing and membrane-free, soluble pyroptolysins was removed, and pellets were resuspended using 1/10 dilutions of detergents stocks from a 96-member Detergent Screen (Hampton). Detergent resuspensions were incubated for 1 hour at 37 °C, centrifuged at 21,000 ×*g* at 4 °C for 30 min with an Eppendorf 5425R mini centrifuge, and the supernatants were analyzed by SDS–PAGE. Select candidate extractions were analyzed by negative-stain electron microscopy and optimized with homemade detergent stocks. The *Lysobacter* pyroptolysin pores were found to extract well with nonylphenyl polyethylene glycol and had an optimal extraction at 4% detergent. The detergent-extracted pores were concentrated by centrifugation, mixed with the amphipol A8-35 at a final concentration of 11 µM, and dialyzed into a buffer containing 20 mM HEPES-KOH (pH 7.5) and 150 mM NaCl at room temperature for 72 hours using 20 kDa MWCO mini dialysis cups (Thermo Fisher Scientific). All extractions were stored at 4 °C until use.

### Cryo-EM sample preparation and data collection

For all pyroptolysin samples, 3 μL of sample was vitrified on grids using a Mark IV Vitrobot (Thermo Fisher Scientific). Thirty min prior to sample vitrification, grids were glow-discharged using an easiGlow (Pelco). Grid type and blotting time was optimized for each sample using a double-sided blot with a constant force of 0, in a 100% relative humidity chamber at 4 °C, and a 10 s wait and a 4–7 s blotting time prior to plunging in liquid ethane before storing in liquid nitrogen. When preparing grids using liposome samples, a ring of parafilm was added to the side of the blot for sample application.

Grids were clipped and first imaged using a Tundra cryo-TEM operating at 100 kV equipped with a Falcon-C direct electron detector (Thermo Fisher Scientific) for initial screening or overnight collections allowing medium-resolution structure determination. EPU software version 3.15.0.11609 was used for all collections. Data for *Lysobacter* pyroptolysin filaments were acquired at a pixel size of 1.14 Å, a total dose of 50.2 e^−^ Å^−2^, and a defocus range of −0.9 to −2.5 µm. Data for pneumolysin filaments were acquired at a pixel size of 1.14 Å, a total dose of 44.1 e^−^ Å^−2^, and a defocus range of −0.9 to −2.5 µm. Cryo-EM data were collected either using a Talos Arctica (Thermo Fisher Scientific) operating at 200 kV or a Titan Krios (Thermo Fisher Scientific) microscope operating at 300 kV. Both microscopes were equipped with a K3 direct electron detector (Gatan). SerialEM software version 3.8.6 was used for all collections. Data collection on grids of the *Lysobacter* pyroptolysin pores on DOPC liposomes with 25 mol% ergosterol were collected using a Talos Arctica (Thermo Fisher Scientific) operating at 200 kV equipped with a K3 direct electron detector (Gatan). SerialEM software version 3.8.6 was used for data collection. Data were acquired at a pixel size of 1.4 Å, a total dose of 181.6 e^−^ Å^−2^ and dose per frame of 1.4 e^−^ Å^−2^. Data collection on grids of the *Lysobacter* pyroptolysin pores extracted from 3:1 DOPC:ergosterol liposomes was performed using a Titan Krios microscope and a total of 11,814 movies were acquired at a pixel size of 0.941 Å, a total dose of 51.7 e^−^ Å^−2^, dose per frame of 0.63 e^−^ Å^−2^ at a defocus range of −0.7 to −2 µm. *Pelomonas* pyroptolysin oligomers and filaments were imaged using a Talos Arctica microscope and a total of 2,656 movies were acquired at a pixel size of 1.1 Å, a total dose of 46.3 e^−^ Å^−2^, dose per frame of 0.99 e^−^ Å^−2^ at a defocus range of −0.6 to −2.0 µm. All micrograph images were prepared using IMOD version 5.1.9.

### Cryo-EM image processing and model building

Movie frames for filaments of *Lysobacter* pyroptolysin (783 movies) and pneumolysin (9,459 movies) were imported to cryoSPARC for patch-based motion correction, patch-based CTF estimation, and 2D classification. Movie frames for the inactive-state oligomers and filaments of *Pelomonas* pyroptolysin were imported to cryoSPARC for patch-based motion correction, patch-based CTF estimation, 2D classification, and non-uniform refinement (see Figure S4 for processing workflow). Movie frames for the *Lysobacter* pyroptolysin pores were imported to cryoSPARC^56^ for patch-based motion correction, patch-based CTF estimation, 2D classification, homogenous refinement, and masked local refinement. Particles were picked using the Topaz^57^ wrapper inside cryoSPARC before extraction and 2D classification (see Figure S7 for processing workflow). Local resolution for both samples was estimated using cryoSPARC and visualized in ChimeraX^55^.

The starting *Pelomonas* pyroptolysin model was obtained from a cryo-EM structure obtained from data collection on a Tundra cryo-TEM (Thermo Fisher Scientific). A model of the active-state *Lysobacter* pyroptolysin generated by ModelAngelo^58^ was fused to an AlphaFold 3^51^ prediction of the Ig-like domain and used as a starting model after docking it into the electron density map of the *Lysobacter* pyroptolysin pore. For all datasets, a complete protomer model was manually built in Isolde version 1.10.1^59^ and iterative corrections were made based on Phenix ^60^ real-space refine. The protomer model was validated in Phenix with MolProbity^61^ (Table S2). The 39-mer pyroptolysin pore model was constructed by extending from a docked protomer by continuously fitting new protomers within the cryo-EM map density. Structure Figure panels were generated using UCSF ChimeraX^55^ and PyMOL^54^ version 2.4.0.

### Structure-based sequence alignments

Structure-based sequence alignments comparing the *Vitiosangium* inactive-state crystal structure and inactive-state *Pelomonas* pyroptolysin cryo-EM structure, the *Vitiosangium* active-state cryo-EM structure and the *Lysobacter* pyroptolysin active-state cryo-EM structure, and crystal and cryo-EM structures of pneumolysin, perfringolysin O, *Lysobacter* pyroptolysin, and *Pelomonas* pyroptolysin were generated using FoldMason^62^. Alignments of desulfolysin and enterolysin sequences with pneumolysin, *Lysobacter* pyroptolysin, and *Pelomonas* pyroptolysin Ig-like domain structures were generated using ProMALS3D^63^ or FoldMason^62^. All alignments were visualized in Jalview version 4.11.5.2^64^.

### Hemolysis assays

Defibrinated sheep or guinea pig blood (Innovative Research, Inc.) was washed in phosphate-buffered saline (PBS, Corning) until the supernatant was clear. Cells were resuspended in PBS to make a 1% erythrocyte solution. Erythrocyte solution and protein were combined in a tissue culture treated 96-well V-bottom plate (Corning 3894) and incubated for 30 mins at 37 °C. The plate was centrifuged for 10 mins at 1,500 *×g* to separate intact erythrocytes from lysate. 50 µL of lysate was transferred to a new plate and heme absorbance was measured at 540 nm using a Synergy HI microplate reader (BioTek). Data was visualized with GraphPad Prism version 10.2.2. The experiment was performed on at least three different occasions using 2–3 technical replicates.

### Liposome assays

Liposomes were constructed from 10 mg of chloroform-suspended lipid stocks of 1,2-dioleoyl-sn-glycero-3-phosphocholine (DOPC), 1-palmitoyl-2-oleoyl-glycero-3-phosphocholine (POPC), or molar ratios of sterol (cholesterol, 3-epicholesterol, ergosterol, β-sitosterol, stigmasterol) and/or sphingomyelin and DOPC or POPC. The epicholesterol was purchased from ALFA Chemistry, while ergosterol, β-sitosterol, and stigmasterol were all purchased from Millipore Sigma. All other lipids were purchased from Avanti. Lipids were dried by transferring to a glass vial with a glass pipette, evaporating chloroform under a stream of nitrogen gas to make a thin lipid film which was held under vacuum overnight in the dark. Liposomes for electron microscopy were prepared by resuspending the dried lipids with 20 mM HEPES-KOH (pH 7.5) and 150 mM NaCl. To prepare liposomes for terbium-based liposome rupture assays, the resuspension buffer was supplemented with 15 mM TbCl_3_ and 50 mM sodium citrate. Liposomes were resuspended in a 1 mL volume (10 mg mL^−1^) by vortexing for 1 min, transferring to a 1.7 mL microfuge tube, and freeze thawing 3× with liquid nitrogen and a warm water bath. The crude liposome suspension was transferred to a glass tube, pulled into a glass syringe (Hamilton), and then extruded (11× for assays and 21× for EM) with a mini-extruder (Avanti) using 19 mm nucleopore membranes with 0.2 µM pores (Whatman) and 10 mm filter supports (Avanti). Extruded liposomes were purified by SEC with a 10/300 Superose 6 column (Cytiva) equilibrated with 20 mM HEPES-KOH (pH 7.5) and 150 mM NaCl. Fractions containing liposomes were combined and stored at 4 °C or −80 °C until use.

Liposome leakage assays were carried out with purified liposomes containing TbCl_3_. 40 µL liposome reactions were assembled in a clear well 96-well PCR plate from 24 µL liposomes and 16 µL dipicolinic acid (DPA) dilution buffer at a final concentration of 20 µM DPA. All assays were initiated by addition of 10 µL of 5 µM protein (for an in-assay concentration of 1 µM) and added to wells containing liposomes. All assays using proteolytic activation were initiated by addition of 5 µL of 10 µM protein, 4 µL of buffer, and 1 µL of 2 mg ml^-1^ proteinase K (NEB) at a 1:1 (w/w) ratio of protease:pyroptolysin. Plate contents were spun to the bottom of wells by brief centrifugation for 1 min at 500 *×g*. The plate was sealed and transferred to a Synergy HI microplate reader (BioTek) to monitor fluorescence resulting from Tb^3+^–DPA complex formation for 6 hours with measurements every min by excitation/emission at 276/545 nm. Data was visualized with GraphPad Prism version 10.2.2. The experiment was performed on at least two different occasions using two technical replicates.

### Bacterial growth spot assays

Pyroptolysin sequences were subcloned into pBAD by PCR amplification from pETS2 templates and Gibson Assembly. pBAD plasmids were transformed into *E. coli* strain DH5α (NEB) and grown in LB media at 37 °C. Whenever applicable, media were supplemented with ampicillin (100 µg mL^-1^) to ensure the maintenance of plasmids. *E. coli* DH5α cells (NEB) containing the pyroptolysin pBAD plasmids were grown overnight in LB medium supplemented with 100 µg mL^-1^ ampicillin and 1% w/v glucose at 37 °C. Saturated cultures were washed with PBS by repeated centrifugation to remove glucose from suppressing arabinose induction. For spot-based growth assays, 10-fold serial dilutions in 1× PBS were performed for each of the samples and 4 µL drops were put on agar plates containing 1% w/v glucose and on plates containing 0.2% w/v arabinose. Plates were incubated overnight at 37 °C. Colony forming unit (CFUs) were determined by counting the derived colonies after overnight incubation. The experiment was performed on at least three different occasions.

### Yeast growth spot assays

Pyroptolysin sequences were amplified from pETS2 and inserted behind a galactose-inducible promoter via Gibson assembly into pRB808 plasmids containing *URA3* (a gift from Kaushik Ragunathan and Ryan Baldrige). Engineered pyroptolysins were then integrated into the *S. cerevisiae* BY4741 genome. Cells were washed in water and normalized to the lowest OD_600_ of any given culture. For the CFU count, 10-fold serial dilutions in water were performed for each of the samples and 2 µL drops were put on YP agar plates containing 2% raffinose and on plates containing 0.2% galactose. Plates were incubated for ∼1 day at 30 °C and 2 days at room temperature before imaging. The experiment was performed on at least three different occasions.

### Homolog searching using Dali

An AlphaFold 3^51^ prediction of *Pseudomonas alcaligenes* ART Rhs-like protein was queried against the full Protein Data Bank using the Dali server^65^. The first 30 amino acids were removed because they were modeled as disordered and had low plDDT scores. The table of hits was downloaded and plotted using GraphPad Prism version 10.2.2.

### Bacterial and yeast liquid culture growth assays

*S. cerevisiae* BY4741, *E. coli* MG1655, *E. coli* DH5α, and *Bacillus subtilis* PY79 strains were used. *S. cerevisiae* cultures were grown in YPD media at 30 °C. *E. coli* and *B. subtilis* cultures were grown in LB media at 37 °C and 30 °C, respectively. Cultures were grown until reaching an OD_600_ of 0.1, after which 100 µL/well was used to inoculate a 96-well plate (GenClone). For protein treatment assays, cells were treated with 5 µM pneumolysin, pyroptolysins, sterile buffer, or antibiotic (kanamycin for *E. coli* and *B. subtilis*) or antifungal (cycloheximide for *S. cerevisiae*). For protein expression toxicity assays, cells were grown to an OD_600_ of 0.5, then diluted in inducing media (LB + 0.25% arabinose) to an OD_600_ of 0.1. Cell growth was monitored over a period of 12 hours with continuous orbital shaking using a Synergy HI microplate reader (BioTek) measuring absorbance at 600 nm every 5 min. Data was visualized with GraphPad Prism version 10.2.2. The experiment was performed on at least three separate occasions with 2–3 technical replicates.

### MTT cytotoxicity assays

All incubation steps were at 37 °C under 5% CO_2_ in an Isotemp CO_2_ incubator (ThermoScientific). Adherent cultures of HEK293 cells were grown to 80–90% confluence in Dulbecco’s Modified Eagle Medium (DMEM, Thermo Fisher Scientific) supplemented with 1X penicillin/streptomycin (Gibco) and 10% fetal bovine serum (Thermo Fisher Scientific). Adherent cells were washed twice with PBS, detached from culture plates (Corning) with trypsin-EDTA (Gibco), and resuspended in DMEM without phenol red (Gibco). Resuspended cells were seeded at a density of 4×10^5^ cells per well in tissue culture treated 96-well V-bottom plates (Corning 3894). Cells were treated with proteins or PBS 24 hours after seeding and incubated overnight. After overnight incubation, 3-(4,5-Dimethyl-2-thiazolyl)-2,5-diphenyl-2H-tetrazolium Bromide (MTT, Millipore) was added to a concentration of 0.5 mg mL^-1^ and incubated for 3.5 hours. Formazan was dissolved by adding an equal volume of 0.1% SDS and 10 mM HCl. 100 µL of the resulting solution was transferred to a 96-well flat bottom plate (Thermo Fisher Scientific) for absorbance measurement at 570 nm using a Synergy HI microplate reader (BioTek) to quantify formazan formation. Data was visualized with GraphPad Prism version 10.2.2. At least three technical replicates were used for each experiment.

### Negative staining electron microscopy

Select pyroptolysins were incubated with DOPC liposomes containing cholesterol or ergosterol at a 3:1 DOPC:sterol molar ratio at room temperature overnight in glass tubes. Reactions were diluted 1/10 in 20 mM HEPES-KOH (pH 7.5) and 150 mM NaCl buffer and 5 µL was applied to glow-discharged 400 mesh copper grids coated with formvar/carbon film (Electron Microscopy Sciences). Grids were blotted after 1 min, stained with 2% uranyl acetate for 2 s, blotted immediately, and air dried. Imaging was performed at 200 kV using a Tecnai F20 TEM using SerialEM Version 4.1.9. Select samples were imaged at 100,000× magnification to acquire enough micrographs for measuring the inner diameters of >50 pores. Pore dimensions were measured manually using ImageJ version 2.9.0. All micrograph images were prepared using IMOD version 5.1.9.

### Comparison of structural transformations between gasdermins, pyroptolysins, and CDCs

An inactive-state model of *Lysobacter* pyroptolysin was generated using Phyre2^66^ with one-to-one threading against the inactive-state structure of *Pelomonas* pyroptolysin. Model visualization and root mean square deviation (RMSD) calculations were done using UCSF ChimeraX^55^. Secondary structure assignments were pulled using the Define Secondary Structure of Proteins (DSSP) implementation^67^ in UCSF ChimeraX^55^ version 1.11. OpenAI ChatGPT (GPT-5.6 Sol; accessed September 13, 2026) was used to assist with development of Python code for quantifying the DSSP secondary structure outputs.

### Sf9 cytotoxicity assays

*Spodoptera frugiperda* (Sf9) suspension cultures were grown in ExpiSF media (Gibco) supplemented with 1× penicillin streptomycin (Gibco) and 1× Fungizone (Gibco) at 27.5 °C. Cells were seeded at a density of 1×10^5^ cells per well in tissue culture treated 96-well V-bottom plates (Corning 3894). Cells were treated with proteins or PBS 24 hours after overnight incubation and incubated for 6 hours before quantifying lactate dehydrogenase release using a CytoTox 96® Non-Radioactive Cytotoxicity Assay kit (Promega). Data was visualized with GraphPad Prism version 10.2.2. At least three technical replicates were used for each experiment.

### Galleria mellonella larvae injections

Endotoxins were removed from purified proteins with Pierce high-capacity endotoxin removal spin columns (Thermo Fisher Scientific) according to manufacturer’s instructions. Protein stocks were aliquoted in sterile tubes, flash frozen in liquid nitrogen, and stored at –80 °C until use. *Galleria mellonella* larvae were purchased from Grubco. Proteins were thawed on ice and diluted in sterile PBS for 10 µL injections using a 25 µl syringe (Hamilton) into a rear proleg of cream-colored larvae without signs of melanization. Larvae were kept in 100 mm petri dishes (Kord-Valmark) at room temperature in the dark and monitored for melanization and response to touch with a pipette tip. Data was visualized with GraphPad Prism version 10.2.2.

---

**Figure S1.**
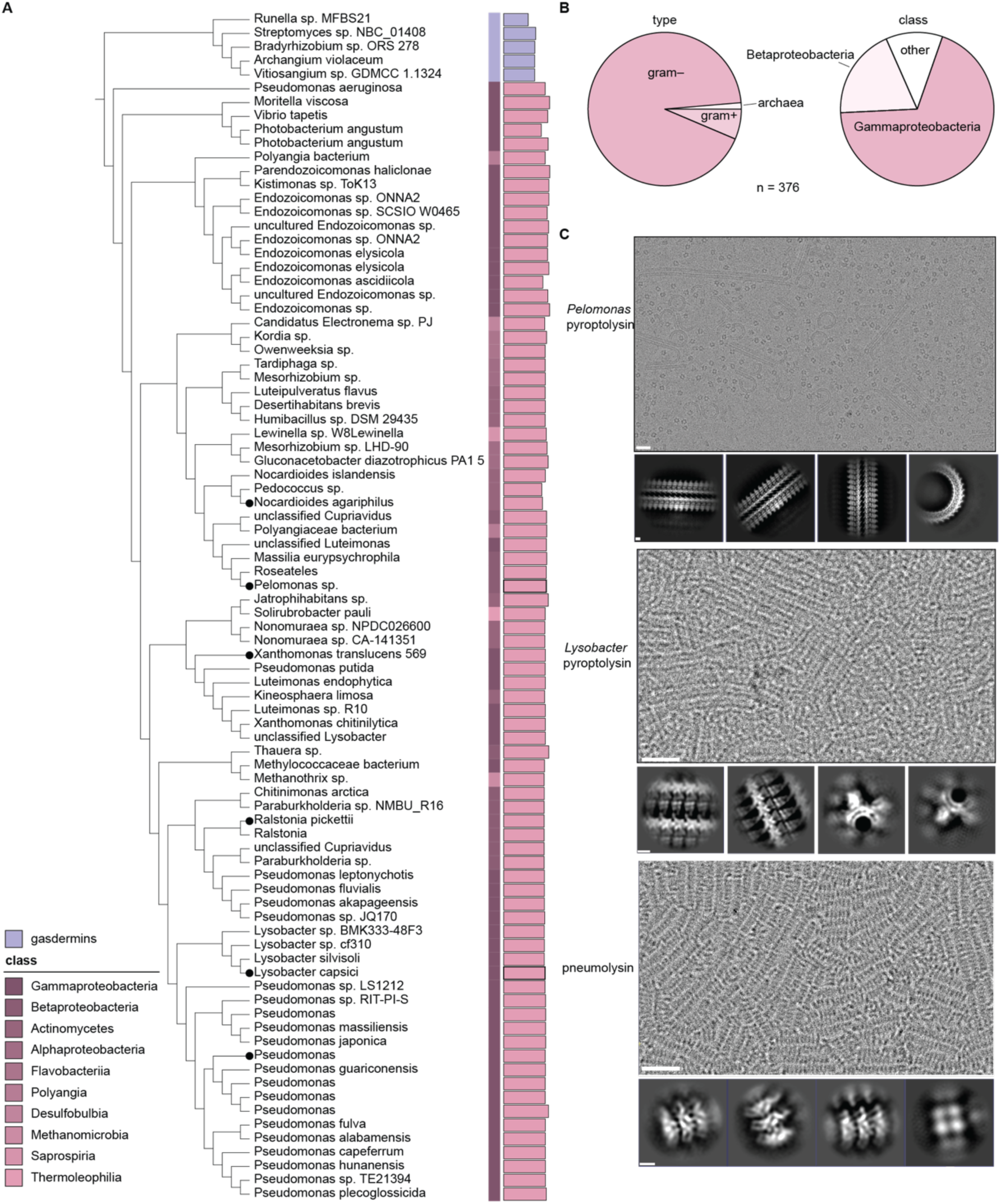
Phylogeny of pyroptolysins. (A) Phylogenetic tree of pyroptolysin protein sequences. Bacterial gasdermins are included as an outgroup. Bar lengths denote sequence length and outer strips are colored by class. (B) Pie charts show species from the nonredundant set of pyroptolysin sequences categorized into major bacterial categories and archaea (left) and taxonomic class (right). (C) Representative cryo-EM micrographs of the *Pelomonas* and *Lysobacter* pyroptolysin filaments, as well as pneumolysin filaments. Scale bars: micrograph, 25 nm; 2D classes, 2.5 nm.

**Figure S2.**
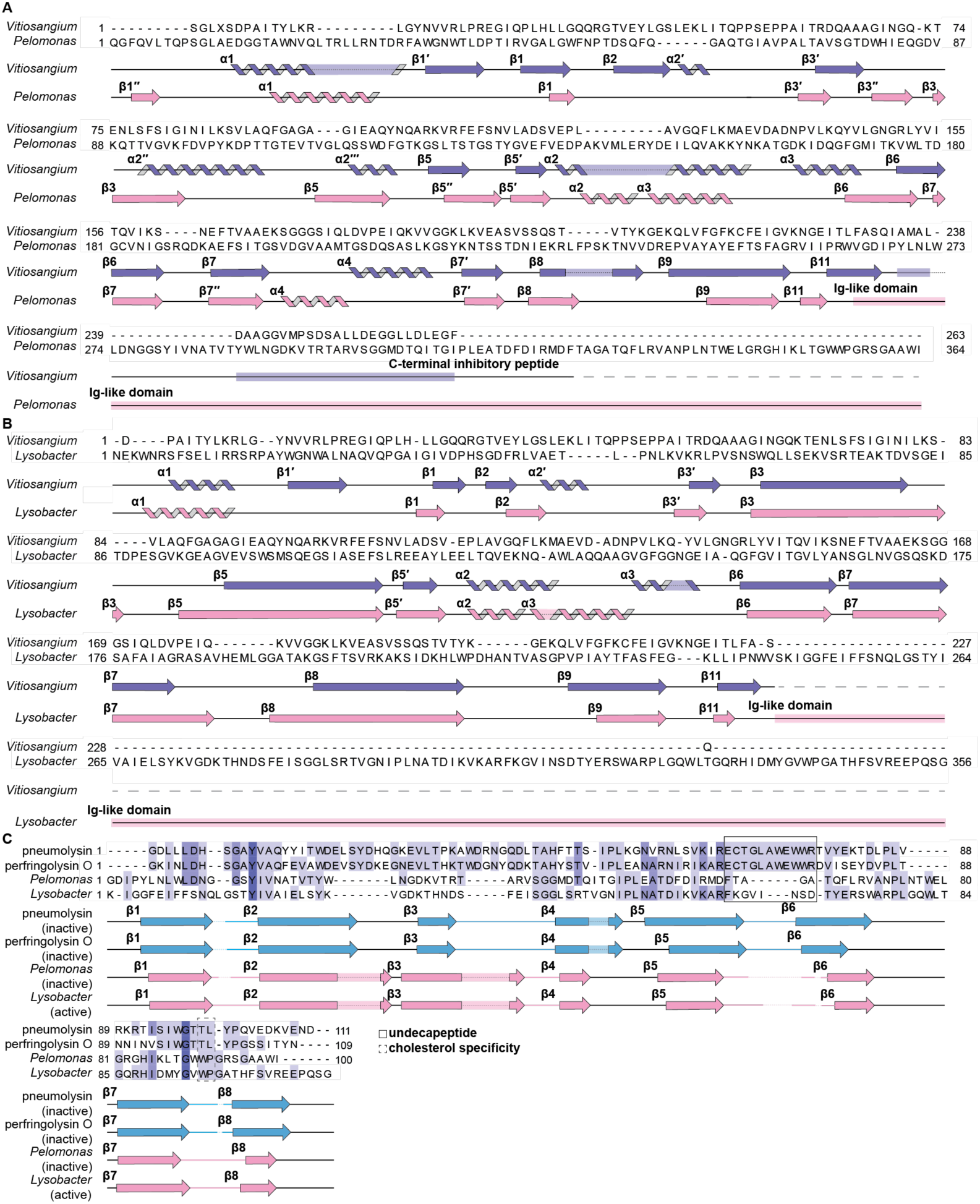
Structure-based sequence alignments of pyroptolysins with gasdermins and cholesterol-dependent cytolysins. (A) Structure-based sequence alignment of inactive-state *Vitiosangium* gasdermin and *Pelomonas* pyroptolysin. Secondary structures are indicated below the sequence alignment for each structure. (B) Structure-based sequence alignment of active-state *Vitiosangium* gasdermin and *Lysobacter* pyroptolysin. Secondary structures are indicated below the sequence alignment for each protein. (C) Structure-based sequence alignment of Ig-like domains from inactive-state pneumolysin and inactive-state perfringolysin O with inactive-state *Pelomonas* pyroptolysin and active-state *Lysobacter* pyroptolysin. Residues are highlighted according to sequence identity, and secondary structure features are indicated below the alignment for each sequence. Secondary structures are indicated below the sequence alignment for each structure. The solid and dashed boxes outline the undecapeptide and cholesterol specificity residues in cholesterol-dependent cytolysins, respectively.

**Figure S3.**
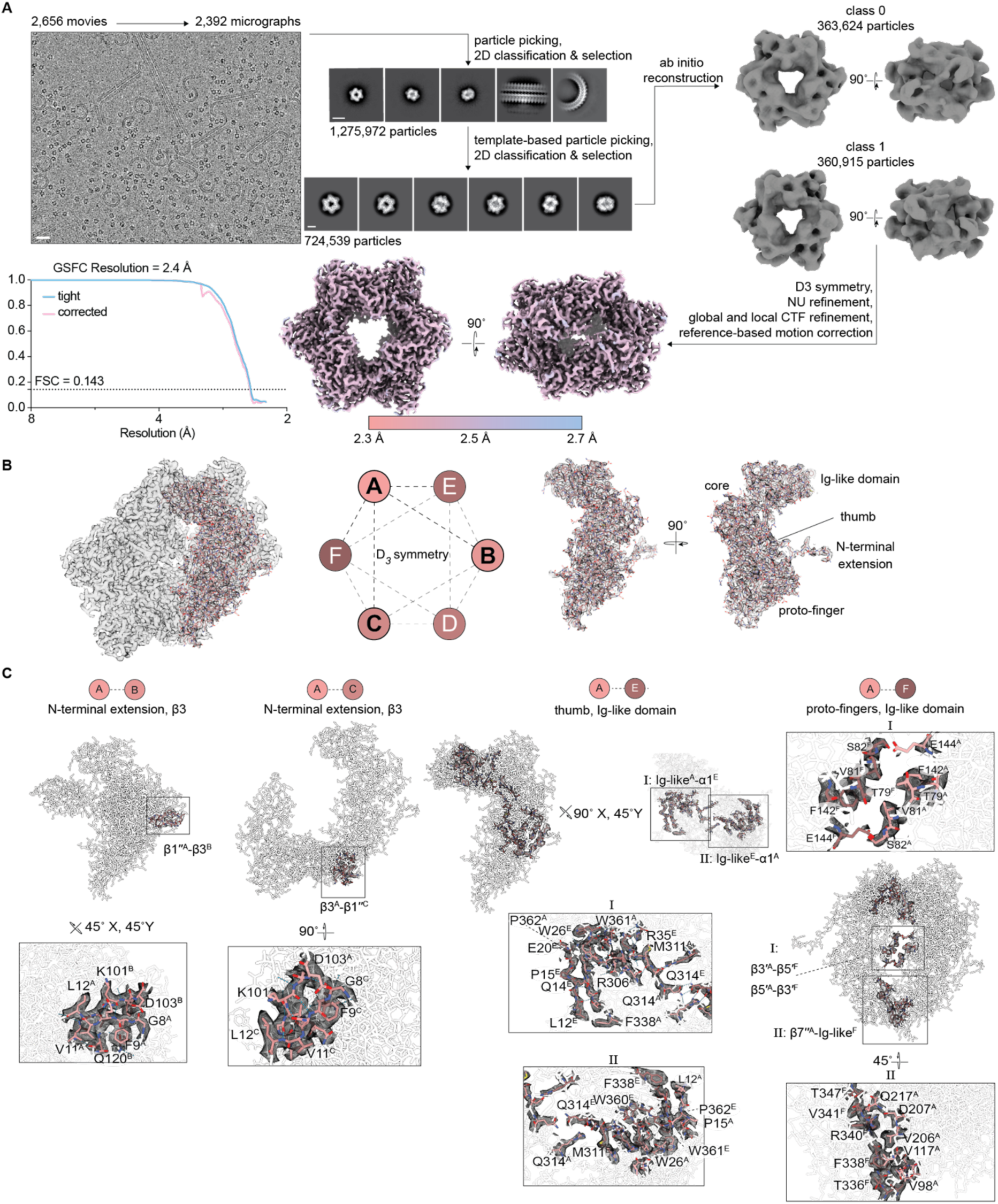
Cryo-EM data processing and map-model overlays of an inactive-state pyroptolysin. (A) Data processing scheme for *Pelomonas* pyroptolysin. Micrograph scale bar, 10 nm; 2D classes scale bar (top), 10 nm; 2D classes scale bar (bottom), 5 nm. (B) Map-model overlays of *Pelomonas* pyroptolysin contoured to 6*σ*. Major structural regions are labeled. Inter-subunit interactions are shown. (C) Map-model overlays of interprotomer interfaces between *Pelomonas* pyroptolysin subunits. Secondary structures are labeled according to gasdermin nomenclature.

**Figure S4.**
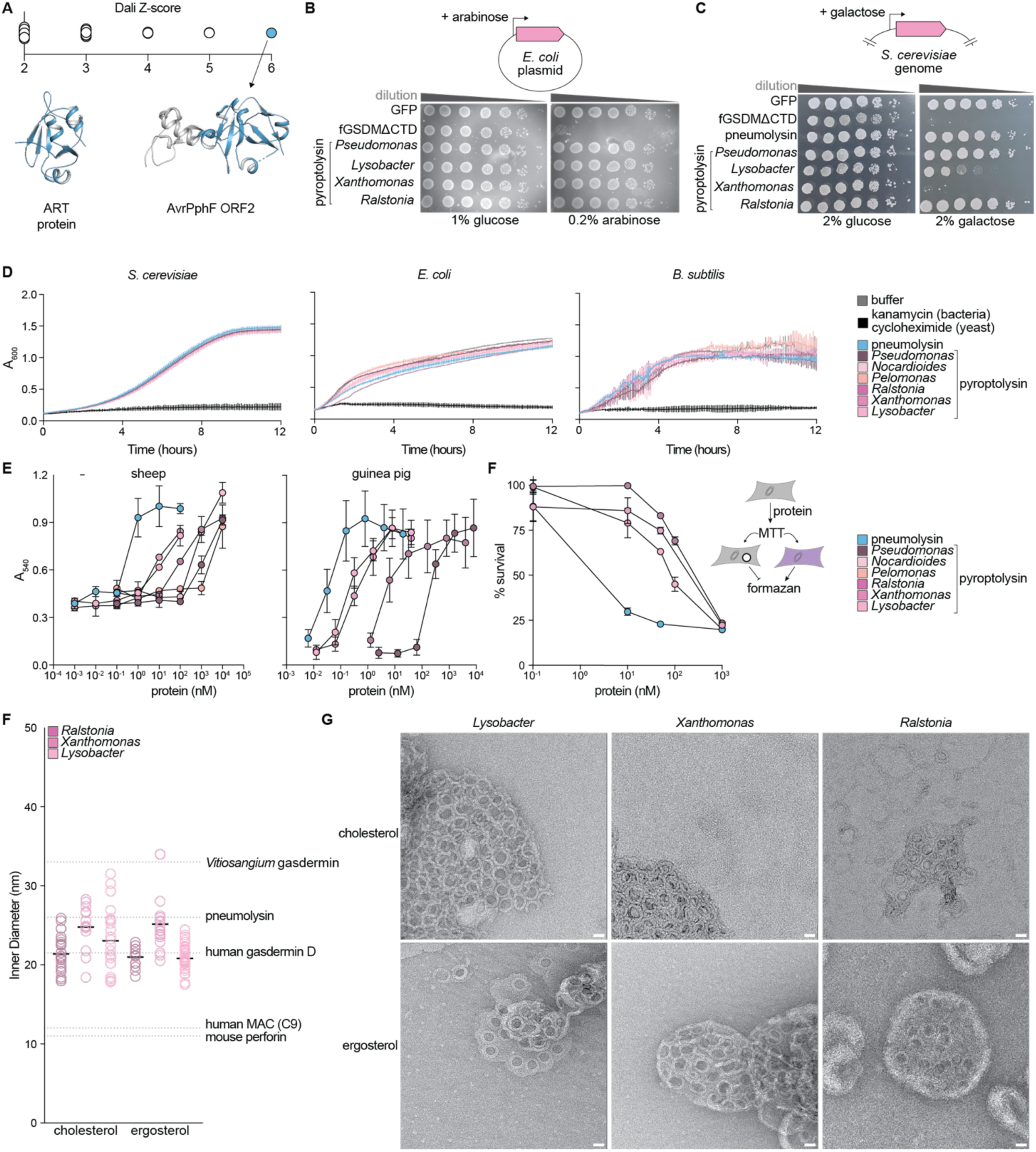
Pyroptolysins have weak activity against mammalian and bacterial cells but can kill yeast cells from the inside and form pores on sterol-containing membranes *in vitro*. (A) Plot of Dali Z-scores of the putative ART protein encoded with the *Pseudomonas alcaligenes* pyroptolysin (see Figure 2). The top hit, AvrPphF ORF2 (Protein Data Bank ID 1S21) is colored blue and an arrow points to the crystal structure of AvrPphF ORF2, beside an Alphafold 3 prediction of the Pseudomonas alcaligenes ART Rhs-like domain. (B) Spot assay of DH5α E. coli harboring plasmids encoding GFP, the autoactive pore-forming domain of a fungal gasdermin from a *Thozetella sp.*, or pyroptolysins behind an inducible promoter. Cells were serially diluted on plates from left (10^0^) to right (10^-6^) with glucose or arabinose to suppress or reduce expression, respectively. (C) Spot assay of *S. cerevisiae* BY4741 harboring genes encoding GFP, the pore-forming domain of a fungal gasdermin from a *Thozetella sp.*, pneumolysin, or pyroptolysins behind an inducible promoter. Cells were serially diluted on plates from left (10^0^) to right (10^-6^) supplemented with glucose or galactose to suppress or reduce expression, respectively. (D) Growth assay of *S. cerevisiae* BY4741, *E. coli* MG1655, or *B. subtilis* PY79 cultures treated with pneumolysin, pyroptolysins, buffer (untreated), or kanamycin (for bacteria) or cycloheximide (for yeast). (E) Dose-dependent hemolysis of guinea pig (left) and sheep (right) erythrocytes after 30 minutes of incubation at 37 °C with pneumolysin and pyroptolysins. (F) Dose-dependent cytotoxicity measured by MTT assay of HEK293 cells treated with pneumolysin and pyroptolysins. (G) Inner diameters of *Lysobacter*, *Xanthomonas*, and *Ralstonia* pyroptolysin pores in DOPC liposomes reconstituted with 25 mol% of cholesterol or ergosterol. Inner diameters of *Vitiosangium* gasdermin (Protein Data Bank ID 9A84), pneumolysin (Protein Data Bank ID 5LY6), human gasdermin D (Protein Data Bank ID 6VFE), human poly-C9 of MAC (Protein Data Bank ID 5FMW) and mouse perforin (Protein Data Bank 7PAG) are indicated as gray lines. (H) Representative negative stain EM micrographs used to measure pore diameter sizes in panel A. Scale bar, 25 nm.

**Figure S5.**
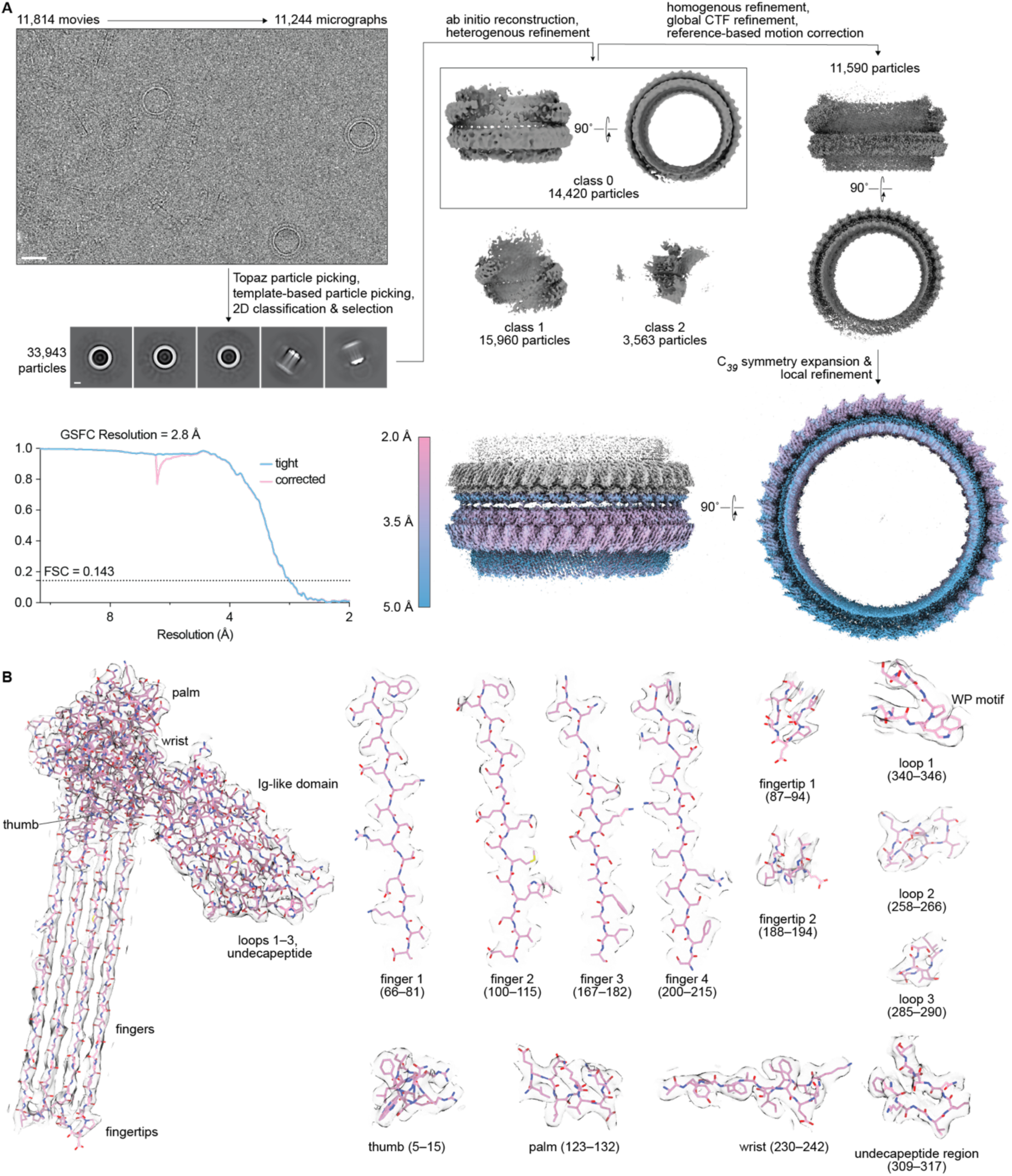
Cryo-EM data processing and map-model overlays of a pyroptolysin membrane pore. (A) Representative cryo-EM micrograph of the *Lysobacter* pyroptolysin pore, 2D class averages from processing micrographs, and single-particle processing schematic. Micrograph scale bar, 25 nm; 2D class scale bar, 5 nm. (B) Example regions of map-model fit quality for a single *Lysobacter* pore protomer. The map surface has been contoured to 6σ.

**Figure S6.**
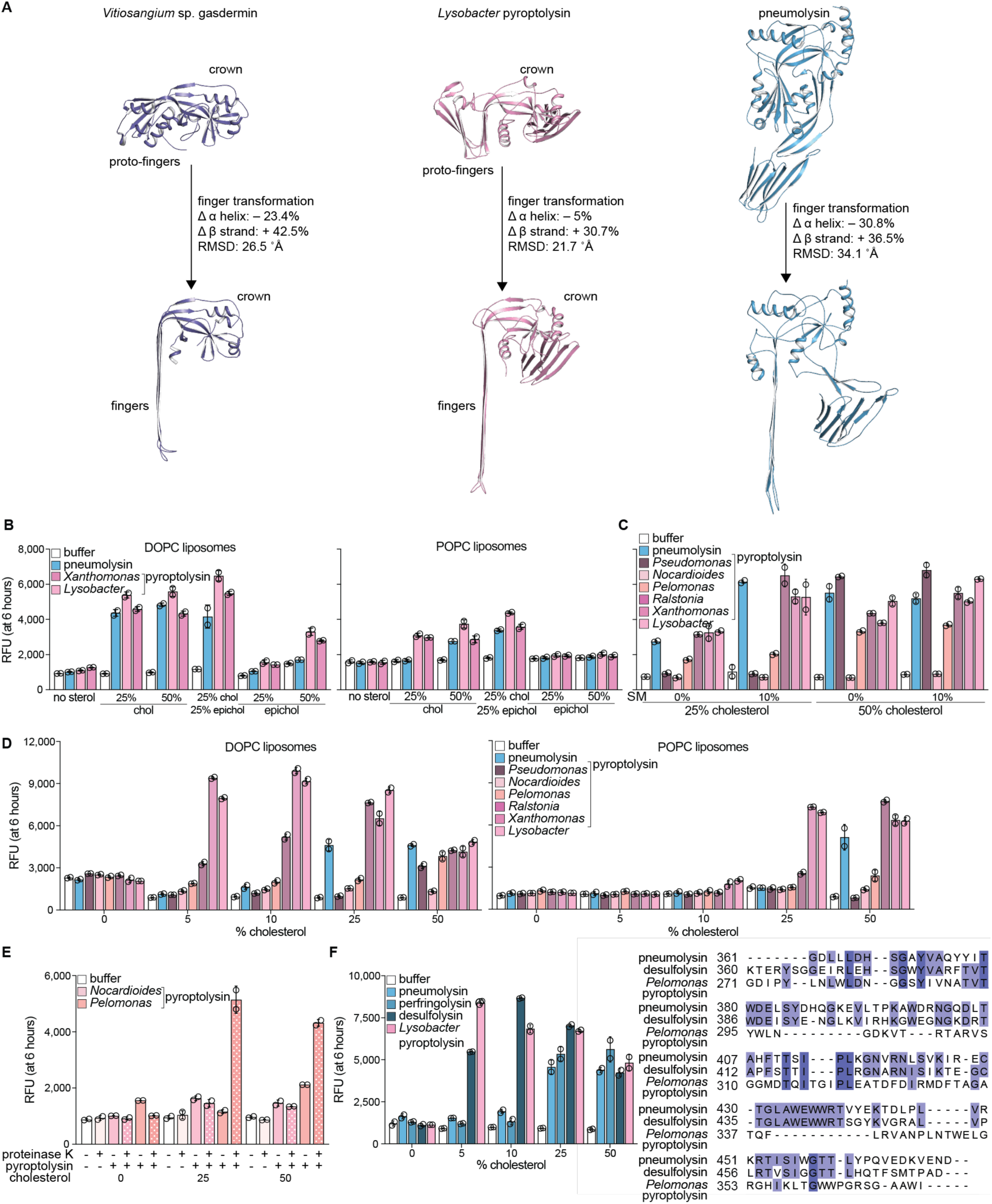
Pyroptolysins are poised for activation and have exceptional sensitivity for accessible membrane sterols. (A) Models of inactive and active-state *Vitiosangium* gasdermin (left, Protein Data Bank IDs 7N51 and 8SL0), inactive and active-state *Lysobacter* pyroptolysin (middle, Protein Data Bank ID 38ON), and inactive and active-state pneumolysin (right, Protein Data Bank IDs 5aod and 5LY6). For each pore-forming protein, the change in α-helix, β-strand, and RMSD between the inactive to active transitions are shown. (B) Liposome rupture experiment testing pneumolysin and pyroptolysins against different base lipids POPC and DOPC with varying concentrations of cholesterol and/or epicholesterol. (C) Liposome rupture experiment testing pneumolysin and pyroptolysins against DOPC liposomes containing different concentrations of cholesterol and sphingomyelin (SM). (D) Liposome rupture experiment testing pneumolysin and pyroptolysins on DOPC or POPC liposomes reconstituted with 5, 10, 25, or 50 mol% cholesterol. (E) Liposome rupture experiment testing *Pelomonas* and *Nocardioides* pyroptolysins with and without addition of proteinase K on DOPC liposomes reconstituted with 0, 25, or 50 mol% cholesterol. (F) Left, liposome rupture experiment testing CDCs and Lysobacter pyroptolysin against DOPC liposomes reconstituted with 5, 10, 25, or 50 mol% cholesterol. For panels A–E, RFUs represent relative fluorescent units at six hours following protein addition. Right, structure-based sequence alignment of the Ig-like domains of inactive-state pneumolysin, an AlphaFold3 prediction of the gram-negative CDC desulfolysin, and inactive-state *Pelomonas* pyroptolysin.

**Figure S7.**
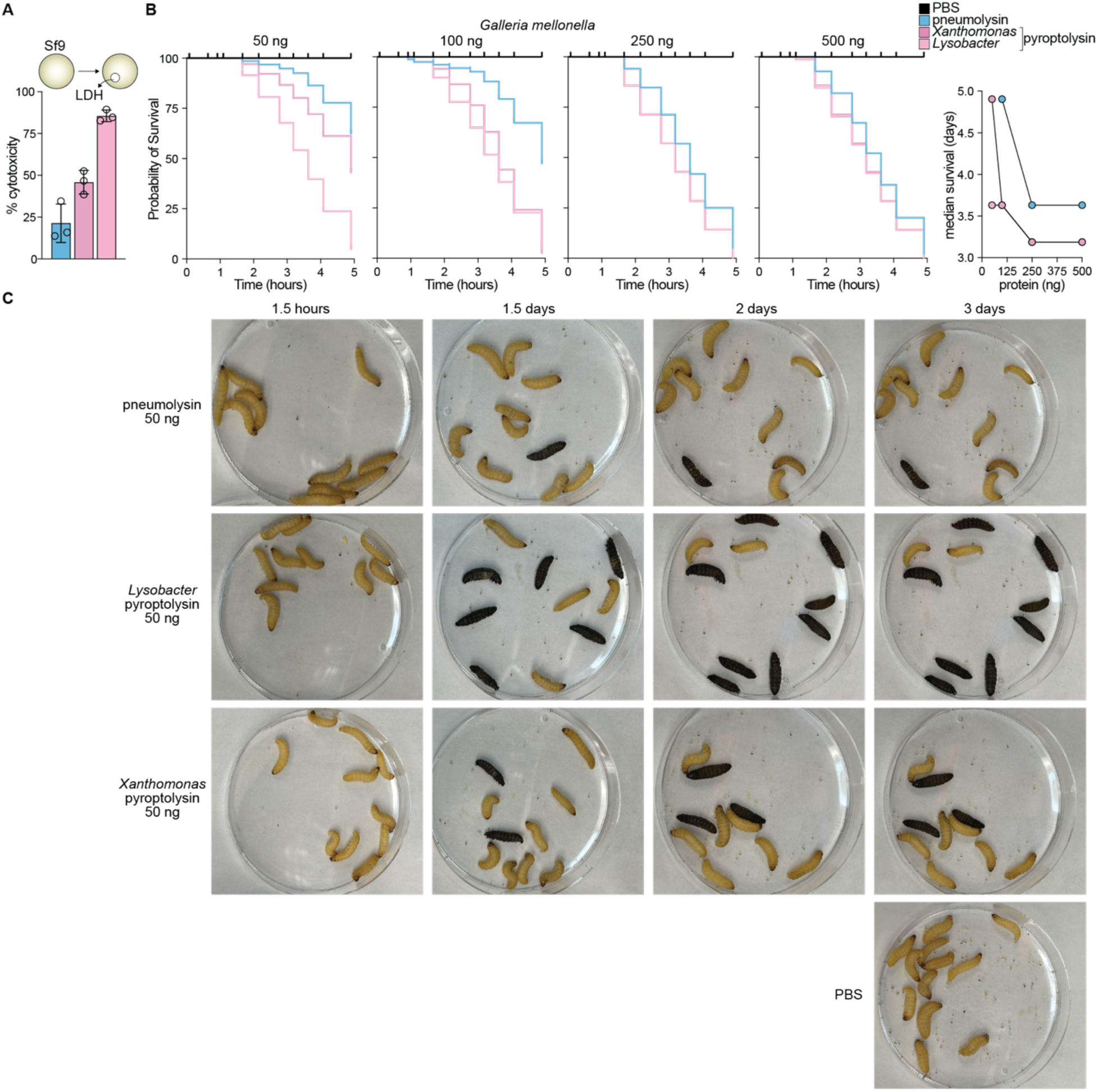
Pyroptolysins have insecticidal activity. (A) Lactate dehydrogenase (LDH) release assay of Sf9 cells treated with 100 nM pneumolysin or pyroptolysins. (B) Left, Kaplan-Meier survival curves of *Galleria mellonella* injected with pneumolysin or pyroptolysins. Right, dose-dependence of the median time of survival. (C) Images of wax moth larvae from selected timepoints after injection with 50 ng of pneumolysin or pyroptolysin.

**Figure S8.**
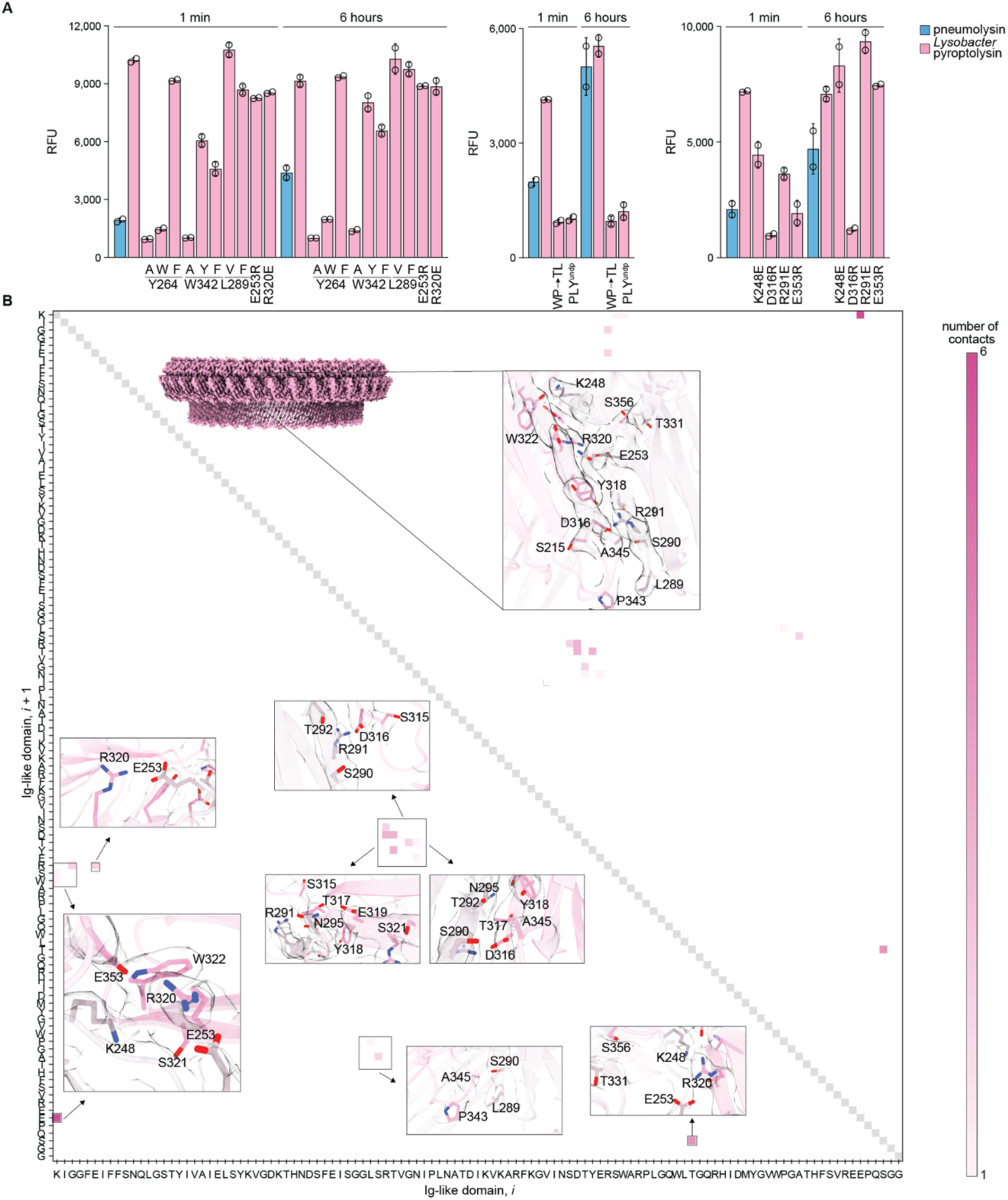
The Ig-like domains of the Lysobacter pyroptolysin pore form an extensive interface. (A) Liposome rupture experiment testing pneumolysin and Lysobacter pyroptolysin mutants against DOPC liposomes with or without 25 mol% cholesterol. RFUs represent relative fluorescent units at five minutes or six hours following protein addition. (B) Heat map of *Lysobacter* pyroptolysin Ig-like domain intersubunit contacts. Insets depict map-model overlays of the Lysobacter pyroptolysin pore corresponding to boxed regions on the heatmap. Adjacent protomers are colored pink and light pink, respectively.

**Table S1.**
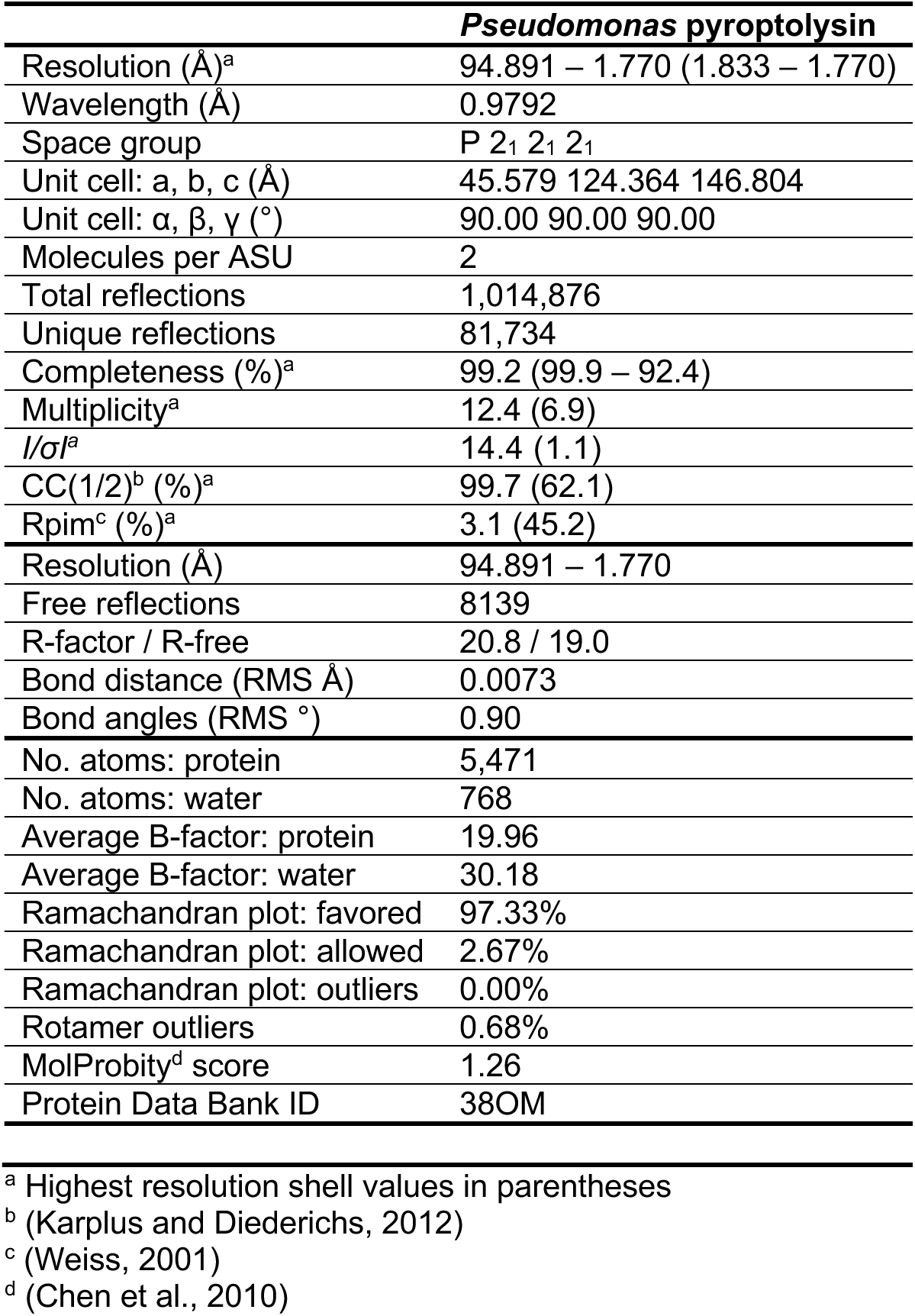
Crystallographic statistics.

**Table S2.** Cryogenic-electron microscopy data summary table.

**Cryo-EM data collection, refinement and validation statistics**
|  | *Pelomonas* pyroptolysin hexamer<br>(EMD-79014)<br>(PDB 38OO) | *Lysobacter* pyroptolysin<br>pore<br>(EMD-79013)<br>(PDB 38ON) |
| --- | --- | --- |
| **Data collection and processing** |  |  |
| Magnification | 36,000 | 130,000 |
| Voltage (kV) | 200 | 300 |
| Electron exposure (e<sup>-</sup>/Å<sup>2</sup>) | 46.2 | 51.7 |
| Exposure time | 5.6 | 4.8 |
| Number of fractions | 40 | 40 |
| Pixel size (Å) | 1.1 | 0.941 |
| Detector | Gatan K3 | Gatan K3 |
| Dose rate detector (e<sup>-</sup>/pix/s) | 8.3 | 9.5 |
| Number of movies | 2,656 | 11,814 |
| Defocus range (μm) | -0.7 to -2.0 | -0.6 to -2.0 |
| Initial particle images (no.) | 724,539 | 84,962 |
| Final particle images (no.) | 360,505 | 11,590 |
| Symmetry imposed | *D*<sub>3</sub> | *C*<sub>39</sub> |
| Map resolution (Å) | 2.4 | 2.8 |
| FSC threshold | 0.143 | 0.143 |
| Map resolution range (Å) | 2.44–3.05 | 2.81–2.09 |
| **Refinement** |  |  |
| Initial model used | ModelAngelo prediction | ModelAngelo prediction |
| Sequences | IMG: 2643742627 | IMG: 8123842218 |
| Model resolution (Å) | 2.1 | 2.8 |
| FSC threshold | 0.143 | 0.143 |
| Model resolution range (Å) | 2.4–2.6 | 2.7–3.2 |
| Map sharpening *B* factor (Å<sup>2</sup>) | -117.5 | -48.89 |
| Model composition |  |  |
| Non-hydrogen atoms | 2,808 | 2,731 |
| Protein residues | 364 | 356 |
| *B* factors (Å<sup>2</sup>) |  |  |
| Protein | 56.72–138.28 | 54.03–209.57 |
| R.m.s. deviations |  |  |
| Bond lengths (Å) | 0.002 | 0.004 |
| Bond angles (°) | 0.464 | 0.973 |
| Validation |  |  |
| MolProbity score | 0.94 | 0.97 |
| Clashscore | 1.81 | 2.04 |
| Poor rotamers (%) | 0.34 | 0 |
| Ramachandran plot |  |  |
| Favored (%) | 99.45 | 98.02 |
| Allowed (%) | 0.55 | 1.98 |
| Disallowed (%) | 0.00 | 0 |

**Table S3.** Comparison of RMSD and DSSP for inactive to active transformations of gasdermins, pyroptolysins, and CDCs.

| **RMSD** |  |  |  |  |  |  |
| --- | --- | --- | --- | --- | --- | --- |
| | Cα RMSD<br>(global) | Cα RMSD<br>(crown) | Cα RMSD<br>(fingers) | # contacts<br>(global) | # contacts<br>(crown) | # contacts<br>(fingers) |
| *Vitiosangium*<br>bGSDM | 25.21 | 3.10 | 26.452 | -64 | 11 | -61 |
| *Lysobacter*<br>pyroptolysin | 31.96 | 2.70 | 21.66 | 370 | 332 | 83 |
| pneumolysin | 23.06 | 3.49 | 34.06 | 12 | 134 | -61 |
| **DSSP (global)** |  |  |  |  |  |  |
| | α-helix<br>(inactive) | α-helix<br>(active) | β-strand<br>(inactive) | β-strand<br>(active) | other<br>(inactive) | other (active) |
| *Vitiosangium*<br>bGSDM | 46 (19.7%) | 16 (7%) | 80 (34.2%) | 117 (51.3%) | 108 (26.2%) | 95 (41.7%) |
| *Lysobacter*<br>pyroptolysin | 25 (10.1%) | 15 (6.1%) | 104 (42.1%) | 135 (54.7%) | 118 (47.8%) | 97 (39.3%) |
| pneumolysin | 80 (22.2%) | 59 (16.4%) | 127 (35.3%) | 115 (31.9%) | 153 (42.5%) | 186 (51.7%) |
| **DSSP (crown)** |  |  |  |  |  |  |
| | α-helix<br>(inactive) | α-helix<br>(active) | β-strand<br>(inactive) | β-strand<br>(active) | other<br>(inactive) | other (active) |
| *Vitiosangium*<br>bGSDM | 24 (17.1%) | 16 (11.9%) | 51 (36.4%) | 48 (35.8%) | 65 (46.4%) | 70 (52.2%) |
| *Lysobacter*<br>pyroptolysin | 20 (13.8%) | 15 (10.3%) | 47 (32.4%) | 47 (32.4%) | 78 (53.8%) | 83 (57.2%) |
| pneumolysin | 33 (19.9%) | 34 (20.5%) | 46 (27.7%) | 22 (13.3%) | 87 (52.4%) | 110 (66.3%) |
| **DSSP (fingers)** |  |  |  |  |  |  |
| | α-helix<br>(inactive) | α-helix<br>(active) | β-strand<br>(inactive) | β-strand<br>(active) | other<br>(inactive) | other (active) |
| *Vitiosangium*<br>bGSDM | 22 (23.4%) | 0 (0%) | 29 (30.9%) | 69 (73.4%) | 43 (45.7%) | 25 (26.6%) |
| *Lysobacter*<br>pyroptolysin | 5 (5%) | 0 (0%) | 57 (56.4%) | 88 (87.1%) | 39 (38.6%) | 13 (12.9%) |
| pneumolysin | 32 (30.8%) | 0 (0%) | 35 (33.7%) | 73 (70.2%) | 37 (35.6%) | 31 (29.8%) |

## Notes

### Summary of Updates

Text and figures were updated, supplemental figures were condensed

## References

1. Shi, J., Zhao, Y., Wang, K., Shi, X., Wang, Y., Huang, H., Zhuang, Y., Cai, T., Wang, F., and Shao, F. (2015). Cleavage of GSDMD by inflammatory caspases determines pyroptotic cell death. Nature 526, 660–665.

2. Kayagaki, N., Stowe, I.B., Lee, B.L., O’Rourke, K., Anderson, K., Warming, S., Cuellar, T., Haley, B., Roose-Girma, M., Phung, Q.T., et al. (2015). Caspase-11 cleaves gasdermin D for non-canonical inflammasome signalling. Nature 526, 666–671.

3. Broz, P., Pelegrín, P., and Shao, F. (2020). The gasdermins, a protein family executing cell death and inflammation. Nat. Rev. Immunol. 20, 143–157.

4. Lieberman, J., Wu, H., and Kagan, J.C. (2019). Gasdermin D activity in inflammation and host defense. Sci. Immunol. 4, eaav1447.

5. Daskalov, A., Mitchell, P.S., Sandstrom, A., Vance, R.E., and Glass, N.L. (2020). Molecular characterization of a fungal gasdermin-like protein. Proc. Natl. Acad. Sci. U. S. A. 117, 18600–18607.

6. Johnson, A.G., Wein, T., Mayer, M.L., Duncan-Lowey, B., Yirmiya, E., Oppenheimer-Shaanan, Y., Amitai, G., Sorek, R., and Kranzusch, P.J. (2022). Bacterial gasdermins reveal an ancient mechanism of cell death. Science 375, 221– 225.

7. Li, Y., Hou, Y., Sun, Q., Zeng, H., Meng, F., Tian, X., He, Q., Shao, F., and Ding, J. (2024). Cleavage-independent activation of ancient eukaryotic gasdermins and structural mechanisms. Science 384, adm9190.

8. Du, G., Healy, L.B., David, L., Walker, C., El-Baba, T.J., Lutomski, C.A., Goh, B., Gu, B., Pi, X., Devant, P., et al. (2024). ROS-dependent S-palmitoylation activates cleaved and intact gasdermin D. Nature 630, 437–446.

9. Rosado, C.J., Kondos, S., Bull, T.E., Kuiper, M.J., Law, R.H.P., Buckle, A.M., Voskoboinik, I., Bird, P.I., Trapani, J.A., Whisstock, J.C., et al. (2008). The MACPF/CDC family of pore-forming toxins. Cell. Microbiol. 10, 1765–1774.

10. Dal Peraro, M., and van der Goot, F.G. (2016). Pore-forming toxins: ancient, but never really out of fashion. Nat. Rev. Microbiol. 14, 77–92.

11. Rosado, C.J., Buckle, A.M., Law, R.H.P., Butcher, R.E., Kan, W.-T., Bird, C.H., Ung, K., Browne, K.A., Baran, K., Bashtannyk-Puhalovich, T.A., et al. (2007). A common fold mediates vertebrate defense and bacterial attack. Science 317, 1548–1551.

12. Ni, T., Jiao, F., Yu, X., Aden, S., Ginger, L., Williams, S.I., Bai, F., Pražák, V., Karia, D., Stansfeld, P., et al. (2020). Structure and mechanism of bactericidal mammalian perforin-2, an ancient agent of innate immunity. Sci. Adv. 6, eaax8286.

13. Ruan, J., Xia, S., Liu, X., Lieberman, J., and Wu, H. (2018). Cryo-EM structure of the gasdermin A3 membrane pore. Nature 557, 62–67.

14. Johnson, A.G., Mayer, M.L., Schaefer, S.L., McNamara-Bordewick, N.K., Hummer, G., and Kranzusch, P.J. (2024). Structure and assembly of a bacterial gasdermin pore. Nature 628, 657–663.

15. Tweten, R.K. (2005). Cholesterol-dependent cytolysins, a family of versatile pore-forming toxins. Infect. Immun. 73, 6199–6209.

16. Gilbert, R.J., Rossjohn, J., Parker, M.W., Tweten, R.K., Morgan, P.J., Mitchell, T.J., Errington, N., Rowe, A.J., Andrew, P.W., and Byron, O. (1998). Self-interaction of pneumolysin, the pore-forming protein toxin of Streptococcus pneumoniae. J. Mol. Biol. 284, 1223–1237.

17. Gilbert, R.J., Jiménez, J.L., Chen, S., Tickle, I.J., Rossjohn, J., Parker, M., Andrew, P.W., and Saibil, H.R. (1999). Two structural transitions in membrane pore formation by pneumolysin, the pore-forming toxin of Streptococcus pneumoniae. Cell 97, 647–655.

18. Peterson, S.B., Bertolli, S.K., and Mougous, J.D. (2020). The central role of interbacterial antagonism in bacterial life. Curr. Biol. 30, R1203–R1214.

19. Bullen, N.P., Sychantha, D., Thang, S.S., Culviner, P.H., Rudzite, M., Ahmad, S., Shah, V.S., Filloux, A., Prehna, G., and Whitney, J.C. (2022). An ADP-ribosyltransferase toxin kills bacterial cells by modifying structured non-coding RNAs. Mol. Cell 82, 3484–3498.e11.

20. González-Magaña, A., Tascón, I., Altuna-Alvarez, J., Queralt-Martín, M., Colautti, J., Velázquez, C., Zabala, M., Rojas-Palomino, J., Cárdenas, M., Alcaraz, A., et al. (2023). Structural and functional insights into the delivery of a bacterial Rhs pore-forming toxin to the membrane. Nat. Commun. 14, 7808.

21. Singer, A.U., Desveaux, D., Betts, L., Chang, J.H., Nimchuk, Z., Grant, S.R., Dangl, J.L., and Sondek, J. (2004). Crystal structures of the type III effector protein AvrPphF and its chaperone reveal residues required for plant pathogenesis. Structure 12, 1669–1681.

22. Farrand, A.J., LaChapelle, S., Hotze, E.M., Johnson, A.E., and Tweten, R.K. (2010). Only two amino acids are essential for cytolytic toxin recognition of cholesterol at the membrane surface. Proc. Natl. Acad. Sci. U. S. A. 107, 4341–4346.

23. van Pee, K., Neuhaus, A., D’Imprima, E., Mills, D.J., Kühlbrandt, W., and Yildiz, Ö. (2017). CryoEM structures of membrane pore and prepore complex reveal cytolytic mechanism of Pneumolysin. Elife 6. 10.7554/elife.23644.

24. Flanagan, J.J., Tweten, R.K., Johnson, A.E., and Heuck, A.P. (2009). Cholesterol exposure at the membrane surface is necessary and sufficient to trigger perfringolysin O binding. Biochemistry 48, 3977–3987.

25. García-Arribas, A.B., Alonso, A., and Goñi, F.M. (2016). Cholesterol interactions with ceramide and sphingomyelin. Chem. Phys. Lipids 199, 26–34.

26. Heuck, A.P., Hotze, E.M., Tweten, R.K., and Johnson, A.E. (2000). Mechanism of membrane insertion of a multimeric beta-barrel protein: perfringolysin O creates a pore using ordered and coupled conformational changes. Mol. Cell 6, 1233–1242.

27. Rosenqvist, E., Michaelsen, T.E., and Vistnes, A.I. (1980). Effect of streptolysin O and digitonin on egg lecithin/cholesterol vesicles. Biochim. Biophys. Acta 600, 91–102.

28. Blumenthal, R., and Habig, W.H. (1984). Mechanism of tetanolysin-induced membrane damage: studies with black lipid membranes. J. Bacteriol. 157, 321–323.

29. Johnstone, B.A., Christie, M.P., Joseph, R., Morton, C.J., Brown, H.G., Hanssen, E., Sanford, T.C., Abrahamsen, H.L., Tweten, R.K., and Parker, M.W. (2025). Structural basis for the pore-forming activity of a complement-like toxin. Sci. Adv. 11, eadt2127.

30. Marheineke, K., Grünewald, S., Christie, W., and Reiländer, H. (1998). Lipid composition of Spodoptera frugiperda (Sf9) and Trichopulsia ni (Tn) insect cells used for baculovirus infection. FEBS Letters 441, 49–52.

31. Vögele, M., Bhaskara, R.M., Mulvihill, E., van Pee, K., Yildiz, Ö., Kühlbrandt, W., Müller, D.J., and Hummer, G. (2019). Membrane perforation by the pore-forming toxin pneumolysin. Proc. Natl. Acad. Sci. U. S. A. 116, 13352–13357.

32. Hotze, E.M., Le, H.M., Sieber, J.R., Bruxvoort, C., McInerney, M.J., and Tweten, R.K. (2013). Identification and characterization of the first cholesterol-dependent cytolysins from Gram-negative bacteria. Infect. Immun. 81, 216– 225.

33. Feil, S.C., Ascher, D.B., Kuiper, M.J., Tweten, R.K., and Parker, M.W. (2014). Structural studies of Streptococcus pyogenes streptolysin O provide insights into the early steps of membrane penetration. J. Mol. Biol. 426, 785–792.

34. Smothers, J.C., Nguyen, C.H.B., Han, Y., Li, Y., Chen, Z., and Radhakrishnan, A. (2026). Visualization of the interaction between sphingomyelin and cholesterol in lipid bilayer membranes. Proc. Natl. Acad. Sci. U. S. A. 123, e2528400123.

35. Johnson, S., Brooks, N.J., Smith, R.A.G., Lea, S.M., and Bubeck, D. (2013). Structural basis for recognition of the pore-forming toxin intermedilysin by human complement receptor CD59. Cell Rep. 3, 1369–1377.

36. Shewell, L.K., Harvey, R.M., Higgins, M.A., Day, C.J., Hartley-Tassell, L.E., Chen, A.Y., Gillen, C.M., James, D.B.A., Alonzo, F., 3rd, Torres, V.J., et al. (2014). The cholesterol-dependent cytolysins pneumolysin and streptolysin O require binding to red blood cell glycans for hemolytic activity. Proc. Natl. Acad. Sci. U. S. A. 111, E5312-20.

37. Shewell, L.K., Day, C.J., Jen, F.E.-C., Haselhorst, T., Atack, J.M., Reijneveld, J.F., Everest-Dass, A., James, D.B.A., Boguslawski, K.M., Brouwer, S., et al. (2020). All major cholesterol-dependent cytolysins use glycans as cellular receptors. Sci. Adv. 6, eaaz4926.

38. Lee, P.-G., Yin, L., Wei, X., Shi, J., Masuyer, G., Wentz, T.G., Chen, P., Xu, Y., Liang, J., Zhang, H., et al. (2025). Identification and characterization of botulinum neurotoxin-like two-component toxins in Paeniclostridium ghonii. Sci. Adv. 11, eadx6145.

39. Xu, Y., Stubbendieck, R.M., Viswanatha, R., Krč, A., Baik, L.S., Suh, W.S., Hu, Y., Wang, H., Yin, L., Mameli, E., et al. (2026). Streptomyces produce a diphtheria toxin-like exotoxin that targets insects. Nat. Microbiol. 11, 1271–1285.

40. Ruhe, Z.C., Low, D.A., and Hayes, C.S. (2020). Polymorphic toxins and their immunity proteins: Diversity, evolution, and mechanisms of delivery. Annu. Rev. Microbiol. 74, 497–520.

41. Das, A., Brown, M.S., Anderson, D.D., Goldstein, J.L., and Radhakrishnan, A. (2014). Three pools of plasma membrane cholesterol and their relation to cholesterol homeostasis. Elife 3. 10.7554/eLife.02882.

42. Jumper, J., Evans, R., Pritzel, A., Green, T., Figurnov, M., Ronneberger, O., Tunyasuvunakool, K., Bates, R., Žídek, A., Potapenko, A., et al. (2021). Highly accurate protein structure prediction with AlphaFold. Nature 596, 583–589.

43. Sayers, E.W., Beck, J., Bolton, E.E., Brister, J.R., Chan, J., Comeau, D.C., Connor, R., DiCuccio, M., Farrell, C.M., Feldgarden, M., et al. (2024). Database resources of the National Center for Biotechnology Information. Nucleic Acids Res. 52, D33–D43.

44. Steinegger, M., and Söding, J. (2017). MMseqs2 enables sensitive protein sequence searching for the analysis of massive data sets. Nat. Biotechnol. 35, 1026–1028.

45. Madeira, F., Madhusoodanan, N., Lee, J., Eusebi, A., Niewielska, A., Tivey, A.R.N., Lopez, R., and Butcher, S. (2024). The EMBL-EBI Job Dispatcher sequence analysis tools framework in 2024. Nucleic Acids Res. 52, W521–W525.

46. Price, M.N., Dehal, P.S., and Arkin, A.P. (2009). FastTree: computing large minimum evolution trees with profiles instead of a distance matrix. Mol. Biol. Evol. 26, 1641–1650.

47. Letunic, I., and Bork, P. (2024). Interactive Tree of Life (iTOL) v6: recent updates to the phylogenetic tree display and annotation tool. Nucleic Acids Res. 52, W78–W82.

48. Cock, P.J.A., Antao, T., Chang, J.T., Chapman, B.A., Cox, C.J., Dalke, A., Friedberg, I., Hamelryck, T., Kauff, F., Wilczynski, B., et al. (2009). Biopython: freely available Python tools for computational molecular biology and bioinformatics. Bioinformatics 25, 1422–1423.

49. Blum, M., Andreeva, A., Florentino, L.C., Chuguransky, S.R., Grego, T., Hobbs, E., Pinto, B.L., Orr, A., Paysan-Lafosse, T., Ponamareva, I., et al. (2025). InterPro: the protein sequence classification resource in 2025. Nucleic Acids Res. 53, D444–D456.

50. Paysan-Lafosse, T., Andreeva, A., Blum, M., Chuguransky, S.R., Grego, T., Pinto, B.L., Salazar, G.A., Bileschi, M.L., Llinares-López, F., Meng-Papaxanthos, L., et al. (2025). The Pfam protein families database: embracing AI/ML. Nucleic Acids Res. 53, D523–D534.

51. Abramson, J., Adler, J., Dunger, J., Evans, R., Green, T., Pritzel, A., Ronneberger, O., Willmore, L., Ballard, A.J., Bambrick, J., et al. (2024). Accurate structure prediction of biomolecular interactions with AlphaFold 3. Nature 630, 493–500.

52. van Kempen, M., Kim, S.S., Tumescheit, C., Mirdita, M., Lee, J., Gilchrist, C.L.M., Söding, J., and Steinegger, M. (2024). Fast and accurate protein structure search with Foldseek. Nat. Biotechnol. 42, 243–246.

53. Tareen, A., and Kinney, J.B. (2020). Logomaker: beautiful sequence logos in Python. Bioinformatics 36, 2272–2274.

54. Schrödinger, L.L.C. (2015). The PyMOL Molecular Graphics System, Version 1.8. Preprint.

55. Pettersen, E.F., Goddard, T.D., Huang, C.C., Meng, E.C., Couch, G.S., Croll, T.I., Morris, J.H., and Ferrin, T.E. (2021). UCSF ChimeraX: Structure visualization for researchers, educators, and developers. Protein Sci. 30, 70–82.

56. Punjani, A., Rubinstein, J.L., Fleet, D.J., and Brubaker, M.A. (2017). cryoSPARC: algorithms for rapid unsupervised cryo-EM structure determination. Nat. Methods 14, 290–296.

57. Bepler, T., Morin, A., Rapp, M., Brasch, J., Shapiro, L., Noble, A.J., and Berger, B. (2019). Positive-unlabeled convolutional neural networks for particle picking in cryo-electron micrographs. Nat. Methods 16, 1153–1160.

58. Jamali, K., Käll, L., Zhang, R., Brown, A., Kimanius, D., and Scheres, S.H.W. (2024). Automated model building and protein identification in cryo-EM maps. Nature 628, 450–457.

59. Croll, T.I. (2018). ISOLDE: a physically realistic environment for model building into low-resolution electron-density maps. Acta Crystallogr. D Struct. Biol. 74, 519–530.

60. Liebschner, D., Afonine, P.V., Baker, M.L., Bunkóczi, G., Chen, V.B., Croll, T.I., Hintze, B., Hung, L.W., Jain, S., McCoy, A.J., et al. (2019). Macromolecular structure determination using X-rays, neutrons and electrons: recent developments in Phenix. Acta Crystallogr. D Struct. Biol. 75, 861–877.

61. Williams, C.J., Headd, J.J., Moriarty, N.W., Prisant, M.G., Videau, L.L., Deis, L.N., Verma, V., Keedy, D.A., Hintze, B.J., Chen, V.B., et al. (2018). MolProbity: More and better reference data for improved all-atom structure validation. Protein Sci. 27, 293–315.

62. Gilchrist, C.L.M., Mirdita, M., and Steinegger, M. (2026). Multiple protein structure alignment at scale with FoldMason. Science 391, 485–488.

63. Pei, J., Kim, B.-H., and Grishin, N.V. (2008). PROMALS3D: a tool for multiple protein sequence and structure alignments. Nucleic Acids Res. 36, 2295–2300.

64. Troshin, P.V., Procter, J.B., and Barton, G.J. (2011). Java bioinformatics analysis web services for multiple sequence alignment--JABAWS:MSA. Bioinformatics 27, 2001–2002.

65. Holm, L., Laiho, A., Törönen, P., and Salgado, M. (2023). DALI shines a light on remote homologs: One hundred discoveries. Protein Sci. 32, e4519.

66. Powell, H.R., Islam, S.A., David, A., and Sternberg, M.J.E. (2025). Phyre2.2: A community resource for template-based protein structure prediction. J. Mol. Biol. 437, 168960.

67. Hekkelman, M.L., Salmoral, D.Á., Perrakis, A., and Joosten, R.P. (2025). DSSP 4: FAIR annotation of protein secondary structure. Protein Sci. 34, e70208.

